# A genome-to-proteome atlas charts natural variants controlling proteome diversity and forecasts their fitness effects

**DOI:** 10.1101/2024.10.18.619054

**Authors:** Christopher M. Jakobson, Johannes Hartl, Pauline Trébulle, Michael Mülleder, Daniel F. Jarosz, Markus Ralser

## Abstract

Despite abundant genomic and phenotypic data across individuals and environments, the functional impact of most mutations on phenotype remains unclear. Here, we bridge this gap by linking genome to proteome in 800 meiotic progeny from an intercross between two closely related *Saccharomyces cerevisiae* isolates adapted to distinct niches. Modest genetic distance between the parents generated remarkable proteomic diversity that was amplified in the progeny and captured by 6,476 genotype-protein associations, over 1,600 of which we resolved to single variants. Proteomic adaptation emerged through the combined action of numerous *cis*- and *trans*-regulatory mutations, a regulatory architecture that was conserved across the species. Notably, *trans*-regulatory variants often arose in proteins not traditionally associated with gene regulation, such as enzymes. Moreover, the proteomic consequences of mutations predicted fitness under various stresses. Our study demonstrates that the collective action of natural genetic variants drives dramatic proteome diversification, with molecular consequences that forecast phenotypic outcomes.

**Highlights:**

- Proteome diversity arises from natural genetic variants, with divergent proteomes in closely related parents and progeny.
- *Cis-*regulatory elements had strong individual impacts, but coherent *trans* effects combined to dominate protein expression.
- Directional selection and frequent transgression suggest much of the proteome is under selective pressure.
- Many *trans*-regulators are enzymes or transporters, with fewer than 4% of pQTLs linking known interactors.
- Genome-to-proteome connections predicted the fitness impact of mutations under various stresses, including a strong but hidden causal variant in *IRA2/*NF1.

## Introduction

Genetic variation plays a central role in health and disease, yet, three decades into the genomic era, we are unable to predict the phenotypic effects of most mutations. For example, the ClinVar database^1^ compiles variants linked to significant clinical effects in well-studied disease genes. Approximately one-third of these variants are classified as being of uncertain significance, and this number continues to grow. The problem is even more acute for rare mutations, which are often presumed to be deleterious but cannot be characterized by population genetics^2^. Nonetheless, accurate functional predictions, if achieved, hold tremendous clinical promise: a study of patients with a monogenic multisystem disease of STAT3, for instance, revealed that all bore mutations causing a biochemical gain-of-function of the protein^3^. These linked challenges arise because we lack a systems-level understanding of how the effects of DNA mutations propagate to other molecular layers and ultimately impact cellular physiology, even in the best-studied organisms. The problem is extremely complex, as mutations may exert their effects on organismal phenotypes by changing the abundance, fold, activity, or otherwise altering the functions and interactions of biomolecules in manifold ways.

Due to rapid progress in nucleic acid sequencing technology, many large-scale efforts to associate mutations with molecular phenotypes have focused on mRNA levels^4^ or mRNA splicing^5^. Yet it is the proteome that predominantly exerts function, and pioneering experiments established the possibility of mapping the effects of variants on protein levels directly^6,7^. This approach has been revolutionized by large-scale antibody-, aptamer- and mass spectrometry-based technologies, primarily focusing on the human plasma proteome^8^. However, two barriers have limited the explanatory power of these datasets. First, the plasma proteome only indirectly represents the primary events of gene expression regulation, being controlled by an interplay of protein excretion by the liver, the tissue leakage of proteins, and glomerular filtration by the kidney. Second, human populations harbor a large excess of rare polymorphisms. As a consequence, genetic associations explain little of the variation observed in plasma protein levels (*e.g.*, 2.7% median genetic contribution in a study with more than 10,000 participants^9^).

On the other hand, a direct link between genetic variation and the proteome can be made in single-cell organisms: the budding yeast *Saccharomyces cerevisiae* is at a sweet spot of genetic tractability due the combination of small genome size and the ability to readily cross and segregate haploid progeny in the laboratory. Crosses of yeast strains have linked genetic variants to changes in mRNA and protein expression at the genome-wide scale^6,10–13^, as well as investigating the regulation of model transcripts and proteins^14–16^. These studies revealed a complex regulatory architecture conserved across eukaryotes, composed of strong *cis*-acting variants alongside pleiotropic *trans*-regulatory mutations (so-called hotspots)^11,17^. Yet the number of proteins or strains examined in proteomic studies of yeast has often been small (∼ 100 segregants)^13,18,19^, and even large collections of wild yeast isolates^20^ are not well-suited to genetic mapping^21^ due to the large number of rare variants. Studies in such panels and in inbred crosses typically cannot resolve linked genomic regions to individual causal polymorphisms, or unambiguously implicate causal genes.

We have shown that this barrier can be overcome by intercrossing the progeny of two closely related wild isolates. Six rounds of meiosis and mating – in contrast to most prior approaches which limited intercrossing to one or two generations – resulted in a panel of haploid segregants in which the genetic linkage between neighboring mutations has been broken, allowing genetic associations to be mapped to individual polymorphisms^22^. Here, we combined precise, systematic proteomics using analytical flow-rate chromatography and Scanning SWATH acquisition^23^ with nucleotide-resolution genetic mapping in a large library of 851 segregants^22^ to comprehensively chart a natural genotype-to-protein map at high resolution. The resulting molecular atlas consisted of thousands of variant-protein associations, many resolved with single-nucleotide resolution and revealed solely at the level of proteins. Notably, the progeny exhibited widespread transgression in proteins not differentially expressed in their ancestors, highlighting the latent potential of the genome to create proteome diversity. Indeed, selection on variants throughout the genome engaged modular regulons to dramatically remodel the proteomes of the two closely related parental strains, revealing general molecular principles underlying causality. Overlaying these molecular data on a complementary genotype-to-phenotype map revealed that the variants controlling protein levels in the absence of stress drove resistance to diverse perturbations. These results suggest that genotype-to-protein maps are conserved across environments and broadly predict phenotypes, charting a path forward to forecast the molecular and phenotypic consequences of genetic variation.

## Results

### Mass spectrometry-based proteomics to probe molecular adaptation

Two ubiquitous obstacles in understanding the mechanistic influence of the genome on the proteome are the excess of rare polymorphisms in natural populations and the difficulty of directly obtaining measurements of protein levels in cells at sufficient scale and precision. Here, we addressed these challenges using 851 F_6_ isolates from a large population of haploid yeast derived from a single mating of two parents, one isolated from the mucosa of an immunocompromised patient (YJM975; henceforth YJM)^24^ and the other isolated from a California vineyard (RM11; henceforth RM)^25^. Despite their substantial phenotypic diversification, they harbor a low level of polymorphism (∼ 0.1%), comparable to that between two unrelated humans. The segregating mutations are in very low linkage disequilibrium, enabling high-resolution genetic mapping^22^.

To measure protein levels in these strains, we took advantage of recent developments in mass spectrometry-based data-independent acquisition (DIA) proteomics using scanning sequential window acquisition of all theoretical mass spectra (Scanning-SWATH)^23^ and new data processing strategies using deep neural networks implemented in the DIA-NN software suite^26^. The high acquisition speed and the ability to match precursor masses with MS2 fragments in Scanning SWATH allowed its integration with high-flow rate analytical chromatography, increasing throughput while maintaining high proteomic depth and excellent quantitative precision. We achieved a measurement throughput of 4.8 min./proteome, compared to, *e.g.*, 120 min./proteome in previous proteome mapping experiments in yeast^13^. We assessed biological and technical variability using numerous controls. The segregant library was cultivated in twelve 96- well plates, each of which included at least three replicates of each parental haploid from which the mapping panel was derived [**Fig. 1A**]. Alongside these, we measured *n* = 117 samples of a pooled sample to detect and correct for batch effects. As a benchmark of species-wide proteome diversity, we also included 22 diverse isolates from the Saccharomyces Genome Resequencing Project (SGRP)^27^ [**Supplemental Table S1**; **Supplemental Table S2**]. We observed low technical variability (C.V. 15.6 - 19.9%) and negligible effects of plate or batch [**Fig. S1ABC**], such that the genetic background was the predominant contribution to proteome variation across the proteomes we acquired [**Fig. 1B**]. The quantified proteins accounted for ∼ 70% of the proteome on a molar basis, and the estimated protein quantities correlated well with absolute protein levels reported previously^28^ [**Fig. S1D**].

**Figure 1.**
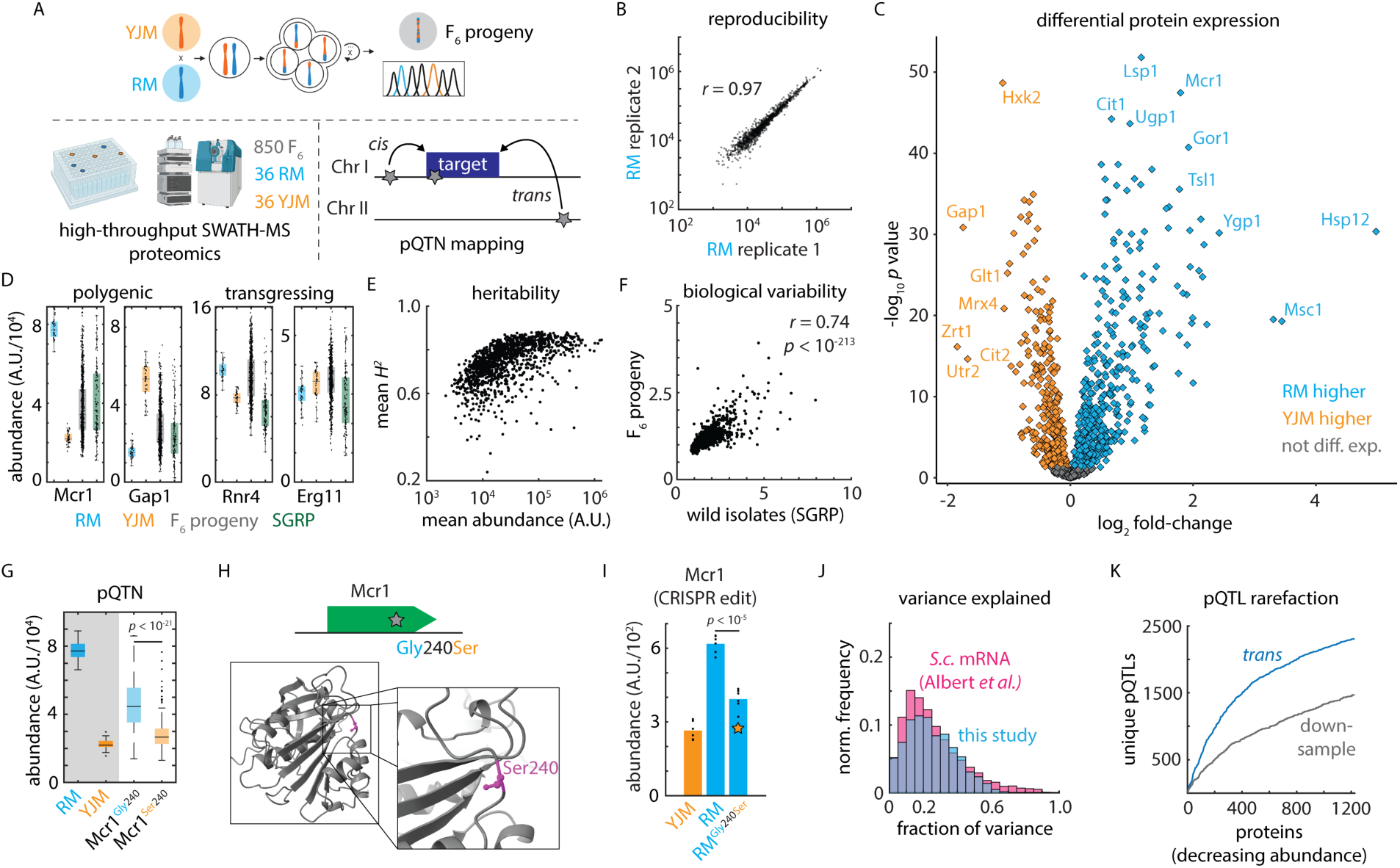
A variant-resolution genome-to-proteome map. (A) Schematic of the mass spectrometry-based proteomics and genetic mapping approach. (B) Representative reproducibility across biological replicates of the vineyard (RM) isolate; Pearson’s *r* as indicated. (C) Volcano plot illustrating log_2_ fold-change in protein abundance (abscissa) and Benjamini-Hochberg-corrected *t* test *p* value (ordinate) between the vineyard (RM) and clinical (YJM) parents. *n* = 36 - 39. (D) Estimated abundance of Mcr1 and Gap1 (polygenic) and Rnr4 and Erg11 (transgressing) in RM parent (blue), YJM parent (orange), F_6_ progeny (grey), and SGRP wild strains (green). Boxes show median and upper and lower quartiles; whiskers show 1.5 times the interquartile range. (E) Mean broad-sense heritability of protein abundance (ordinate) as a function of estimated absolute protein abundance (abscissa) for all proteins measured in at least 80% of samples. (F) Normalized C.V. amongst the SGRP wild strains as compared to the mean C.V. in the parental isolates (ordinate) as a function of normalized C.V. amongst F_6_ progeny (abscissa). Pearson’s *r* as indicated. *p* value by *t* statistic. (G) Genetic mapping of a *cis*-acting SNP controlling the abundance of Mcr1. (H) Schematic and predicted AlphaFold2 protein structure of a *cis*-acting missense variant in Mcr1. (I) CRISPR reconstruction and mass spectrometry to validate the effect of the Mcr1^Gly240Ser^ variant. *n* = 6; *p* value by two-sided *t* test. (J) Histogram of the fraction of total variance explained by the global (*cis*- and *trans*-acting) model in this study (blue) and in a highly powered eQTL mapping study in budding yeast (pink) ^12^. (K) Rarefaction plot of unique *trans-* acting pQTL associations (blue) discovered, ordered by decreasing estimated protein abundance. Also shown in grey is the same statistic for downsampled real data using only 50% of the F_6_ progeny. See also Figure S1.

### Standing and latent variation in the proteome

Despite modest genetic distance (∼ 12,000 mutations; ∼ 1 - 2 x 10^6^ divisions since the last common ancestor^29^) and similar growth properties in standard laboratory conditions [**Fig. S1E**], the proteomes of the parents were highly diverged. For 826 of the 1,225 proteins quantified in the two parents (67.4%), we obtained significantly different intensities (*n* = 36 - 39; B.-H. corrected *q* value < 0.05; 190 with fold-change > 1.5; 66 with fold-change > 2) [**Fig. 1C**]. The most up- and down-regulated subsets of the proteome were highly functionally coherent: for example, the clinical isolate (YJM) had higher levels of amino acid and purine biosynthesis and gluconeogenesis proteins, whereas the vineyard isolate (RM) had higher levels of proteins associated with oxidative phosphorylation and the TCA cycle [**Supplemental Table S3**]. These differences correspond broadly to the two key metabolic states of budding yeast, reflecting a fermentative versus a respiratory metabolism, respectively.

Protein abundance spanned a large dynamic range, both between the parents and amongst the F_6_ progeny, as many protein levels in progeny transgressed beyond their abundance in the parental strains [**Fig. 1D**]. Despite this, our approach yielded very high broad-sense heritability (median 76.2%), which depended only modestly on protein abundance [**Fig. 1E**] and was limited primarily by technical variability rather than gene-by-environment interactions [**Fig. S1F**]. Transgression was common, and, indeed, the variation amongst the progeny was greater than that between the parents for 77.9% proteins we measured (955 of 1,225). Strikingly, the proteomic variation released in the F_6_ progeny was most pronounced for the proteins that were also highly variable across genetically diverse wild yeast isolates spanning the diversity in this species ^27^ (*r* = 0.74; *p* < 10^-213^) [**Fig. 1F**]. Thus, the proteomic diversification released by meiosis in our experiment was broadly representative of species-wide variation, perhaps reflecting conserved layers of modular regulation in this organism.

### A nucleotide-resolution proteogenomic map in a model eukaryotic species

Based on these high-quality measurements and the statistical power afforded by the F_6_ segregant panel, we performed genetic mapping^22^ to identify variants associated with changes in protein abundance. Briefly, we conducted global and *cis*-focused (local) mapping by multivariate regression [**Fig. 1A**], including growth differences as a covariate. This proved important for a small subset of proteins, as found previously^12,30^ [**Fig. S1G**]. The effects of associations that were discovered in both local and global mapping agreed well [**Fig. S1H**].

Global mapping, which encompassed all segregating polymorphisms and allowed us to compare *cis*- and *trans*-acting effects, identified 6,476 variant-protein associations (pQTLs) controlling the abundance of 923 proteins (∼ 10% FDR; by permutation; see **Methods**) [**Supplemental Table S4**]. Of these, 1,650 of the associations (25.5%) fine-mapped to a single underlying polymorphism, granting an unprecedented molecular window onto the genome-to-proteome map. In the case of the mitochondrial NADH-cytochrome *b*5 reductase Mcr1^31^, for example, we identified a coding SNP (Mcr1^Gly240Ser^) that was associated with reduced Mcr1 levels in *cis* [**Fig. 1GH**]. Upon reconstruction of the variant by genome editing, subsequent proteome analysis revealed that the Mcr1^240Ser^ mutation alone was sufficient to decrease Mcr1 level by nearly 40% [**Fig. 1I**].

Our model explained a median of 30.4% of the broad-sense heritability in protein level, and, due to the high heritability of protein abundance in our experiment, we explained 22.8% of the variance in protein abundance [**Fig. 1J**]. This was comparable to mapping of mRNA abundance in yeast (median 21.9% variance explained^12^). Our approach, however, achieved much higher resolution: the median confidence interval in prior studies of yeast crosses ranged from 48 kb (mRNA eQTL^12^) to 68 kb (protein X-pQTL^14^). Moreover, approaches such as X-pQTL mapping that rely on tagged proteins freeze the immediate genomic context, prohibiting direct assessment of *cis*-acting effects. We were well-powered to detect additional associations of modest effect had they been present (sensitivity ∼ 95% for effects of 0.1 standard deviations; ∼ 63% for 0.025 s.d.) [**Fig. S1I**]. Thus, residual missing heritability in our map was likely due to numerous additional pQTLs of small effect or, potentially, epistatic interactions.

As expected given the high sensitivity of our mapping panel, the rate at which we discovered additional unique *trans* pQTLs declined as we considered additional proteins [**Fig. 1K**], suggesting that we captured a comprehensive overall picture of protein regulation. At the same time, downsampling real data to 50% of the strains in the experiment yielded just 3,498 associations (54% of the complete atlas), confirming that we were well-powered to chart the regulatory network. In concordance with widespread transgression, we identified at least one pQTL for 233 of the 399 proteins that were not differentially expressed between the parents (mean 2.63 pQTLs per protein) [**Fig. S1J**]. Accordingly, the true biological variability released in the cross (C.V. amongst the F_6_ progeny normalized to technical C.V.) was highly predictive of the number of pQTLs discovered for a protein (*r* = 0.60; *p* < 10^-117^) [**Fig. S1K**]. Overall, across the ∼1,200 proteins we robustly quantified, at least 1,000 were subject to genetic control (as indicated by differential expression or regulation by a pQTL), even in the closely related isolates we analyzed. Thus, our approach presents an opportunity to understand the molecular genetic basis of both standing and latent variation in the proteome.

### Testing the impact of causal variants across the species

As a test of our mapping findings, we next examined the penetrance of pQTL effects across other natural isolates, exploiting the transcriptomes and proteomes^32^ of the 1,002 Yeast Genomes collection^20^ [**Fig. 2A**]. Across these diverse wild strains, both Odc2 and Rdl1 transcript and protein levels, for example, were affected by the *cis*-acting variants identified [**Fig. 2B**]. Broadly, *cis*- acting variants affected the same protein abundances across the divergent natural strain backgrounds in this independent experiment (Mann-Whitney *U* test *p* < 10^-3^; 46 concordant out of 67 *cis*-pQTL associations tested) [**Fig. S2A**]. Strikingly, we also identified several instances (*e.g.* Faa1 and Map1) in which protein *cis*-regulatory effects were evident at the proteome but not at the transcriptome [**Fig. 2B**]. Thus, our protein-oriented mapping captured both mRNA regulation that propagated to protein levels as well as the molecular basis of regulation that emerged primarily in the proteome^33^, with these effects evident species-wide.

**Figure 2.**
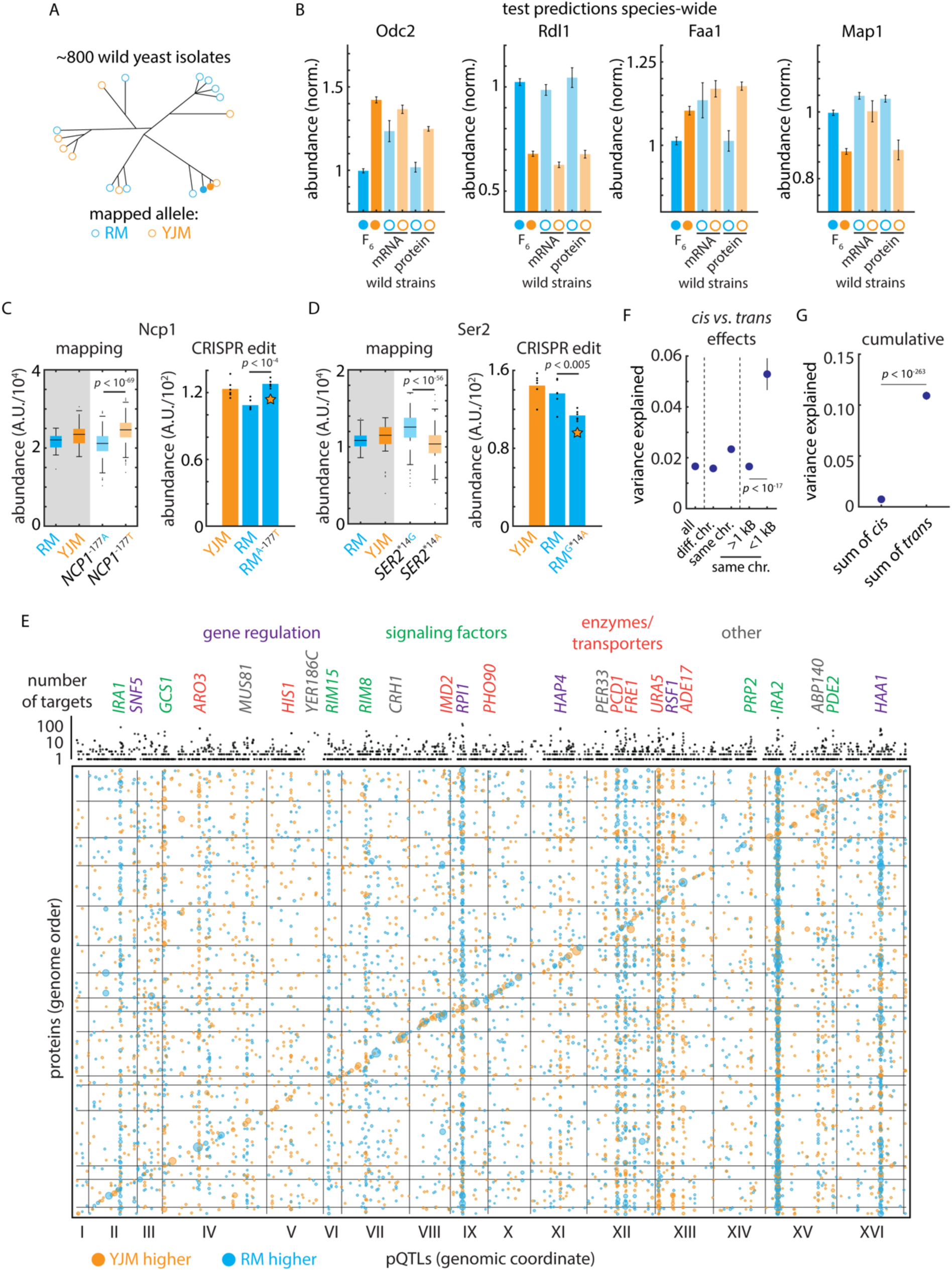
Mutation-to-molecule atlas reveals protein-level regulation. (A) Schematic of statistical replication strategy. (B) Left: Genetic mapping of *cis*-acting effects on Odc2 and Rdl1 protein abundance and replication of this signal in the orthogonal 1,002 Yeast Genomes transcriptomes and proteomes. Right: As left, but for Faa1 and Map1; these signals were evident only at the proteomic level in the replication data. Data shown are median and s.e.m. (C) Left: Genetic mapping of *cis*-acting effect on Ncp1 protein abundance. Right: CRISPR reconstruction and mass spectrometry to test the effect of the *NCP1*^A-177T^ variant. *n* = 6; *p* value by two-sided *t* test. (D) As in (C), but for the *SER2*^G*14A^ variant. (E) Bubble plot indicating the genomic position of all pQTLs. pQTL positions and encoding genes are arranged in genome order. Orange dots indicate clinical (YJM) allele increases protein level; blue indicates vineyard (RM) allele increases level. Dots are sized by genetic mapping *p* value. Indicated above is the number of target proteins controlled by each locus (aggregated by gene); highlighted are *trans* hotspots color-coded by gene function as indicated. (F) Variance explained by pQTLs with the indicated distance to the encoding gene for the target protein; *p* values by Student’s *t* test. Dots indicate mean and bars standard error. (G) Cumulative effect of *cis*- and *trans*-acting pQTLs across all proteins. Dots indicate mean and bars standard error; *p* value by Student’s *t* test. See also Figure S2.

### mRNA- and protein-level gene regulation

We then compared our protein mapping data with allele-specific mRNA expression (ASE) analysis of the F_0_ hybrid of the parents of our genetic mapping panel [**Fig. S2B**]^34^. Interestingly, only 30 of 127 proteins with a *cis*-pQTL had a significant mRNA allelic imbalance, even though we were well-powered to detect allele-specific expression of these mRNAs (117 of the *cis*-pQTLs had a tag SNP in the associated ORF; median depth 183 read counts) [**Supplemental Table S5**]. These data indicate that many *cis* effects arise more strongly at the protein level rather than at the mRNA. This could occur if a variant affects the translation, folding, trafficking, or localization of the encoded protein.

To examine this property in detail, we selected two regulatory *cis*-pQTNs, one mutation upstream of the *NCP1* gene encoding a P450 reductase and one in the 3’ untranslated region (UTR) of *SER2*, which encodes phosphoserine phosphatase. The effect of the *NCP1* mutation was only significant for protein level (no mRNA ASE was detected), while the *SER2* variant impacted mRNA and protein levels in similar fashion. We then used CRISPR genome editing to reconstruct these mutations^35^ and used proteomics to measure protein abundances. In both cases, the exchange of the variant recapitulated the predicted effects: the *NCP1*^A-177T^ mutation resulted in an upregulation of Ncp1 (*p* < 10^-4^), while introducing *SER2*^G*14A^ downregulated Ser2 (*p* < 10^-4^) [**Fig. 2CD**].

### Non-canonical regulators underlying trans-acting hotspots

Our genotype to proteome atlas reflects considerable complexity in the regulation of protein expression: the median protein was controlled by 5 loci and 22.6% of proteins were controlled by more than 10 pQTLs [**Fig. S2C**]. 98% of associations involved distant, presumably *trans*-acting loci (> 1 kB from the target gene in the compact *S. cerevisiae* genome) while the remainder were nearby and likely acted in *cis*. A large proportion of these associations were due to a small number of *trans*-regulatory hotspots^10,11^ that controlled a disproportionate number of targets: the 100 most pleiotropic *trans*-pQTL genes (out of ∼ 2,000) accounted for more than 44% of associations.

The transcription factor *PHO2*, for instance, controlled the adenine biosynthetic pathway [**Fig. S2D**]. Notably, however, many hotspots did not arise from DNA-binding proteins or regulatory factors, but rather metabolic enzymes or membrane transporters [**Fig. 2E**] The uracil transporter *FUR4* controlled the uracil biosynthetic pathway, and the inosine monophosphate dehydrogenase *IMD2*, involved in GTP synthesis, controlled the abundance of a variety of other metabolic enzymes [**Fig. S2D**]. The effects of these highly functionally coherent regulons combined with *cis-*acting variants to produce large changes in protein abundance amongst the haploid progeny. Although nearby *cis-*acting variants were of larger effect (mean 5.29% of variance explained vs. 1.66%, *p* < 10^-17^ by Mann-Whitney *U* test) [**Fig. 2F**], the cumulative effect of *trans* regulation on a typical protein was much larger (mean 10.9% of variance explained in *trans* vs 0.74% in *cis* across all proteins, *p* < 10^-263^ by Mann-Whitney *U* test) [**Fig. 2G**]. This comprehensive atlas positioned us to investigate how natural genetic variation drives proteomic adaptation through the action of multiple *trans*-regulatory hotspots throughout the genome.

### Regulatory adaptation underlying diverged proteomes

Examining the proteins upregulated in the parental isolates, transcription factor target analyses^36^ indicated that the YJM-upregulated gene set was highly enriched for targets of the Sfp1, Stb3, Dot6, Tod6, and Gcn4 transcription factors, whereas the YJM-downregulated module was likely regulated by Sut1, Msn2/4, Hap3/5, and Gis1 [**Supplemental Table S6**]. Yet there were no *trans*- regulatory hotspots at the genes encoding these factors. We therefore scrutinized our genotype-to-protein map further to identify other possible origins of these proteomic changes. We found that three of the most pleiotropic *trans*-acting loci in our experiment (*IRA1*, *IRA2*, and *PDE2*) were centered at genes in the Ras/PKA pathway^37–39^, a signaling pathway conserved from yeast to humans^40^. The Ras/PKA network integrates nutritional signals to control metabolism and proliferation and is associated with adaptation to fermentation^41^ as well as virulence in pathogenic yeasts^42^. Two well-characterized targets of the Ras/PKA signaling pathway (*via* the kinase Rim15^43^) are the Gis1 and Msn2/4 transcription factors, consistent with our transcription factor target analyses.

The three hotspots at *IRA1*, *IRA2*, and *PDE2* [**Fig. 3A**] controlled the abundance of 50 to over 300 proteins, with coherent subsets of proteins up- and down-regulated by each parental allele. The abundance of Mcr1, for example, ranged nearly 3-fold depending on the genotype at just 3 hotspot loci and a single *cis*-acting SNP at the *MCR1* locus [**Fig. 3B**]. To visualize the concerted effects of these alleles, we generated a *t*-distributed stochastic neighbor embedding (*t*- SNE) of the correlations in protein abundance. Proteins that were significantly upregulated in the clinical and vineyard strains formed pronounced clusters, and we noted that a similar set of proteins was differentially regulated by each Ras/PKA hotspot [**Fig. 3C**]. Consistent with our hypothesis that these variants controlled downstream transcriptional activation, our genetic mapping results agreed well with the effects of *IRA1*, *IRA2*, and *PDE2* deletions on transcript abundance [**Fig. S3A**]^44^.

**Figure 3.**
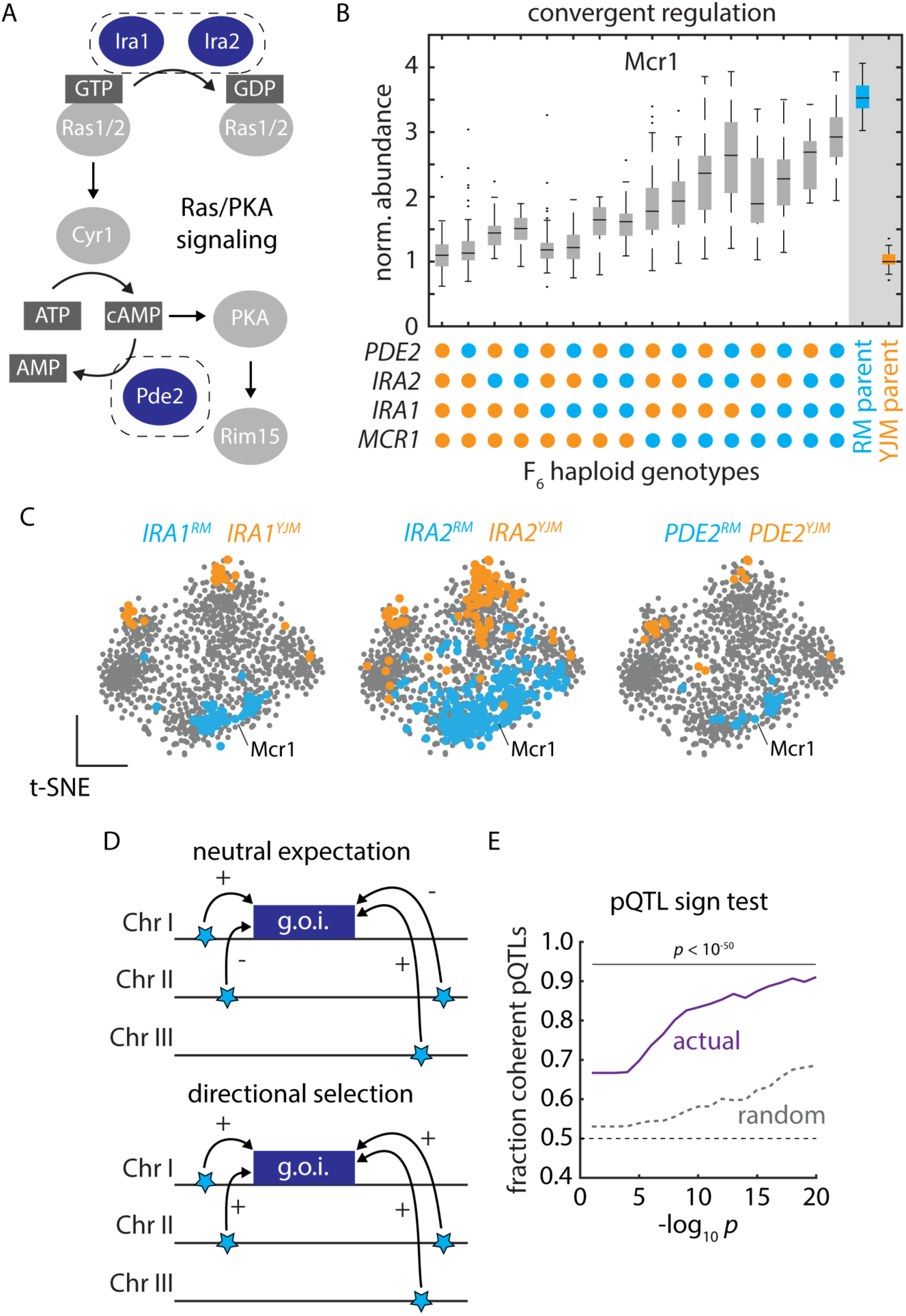
Polygenic adaptation reflecting natural selection on protein abundance. (A) Schematic of Ras/PKA signaling highlighting the Ira1, Ira2, and Pde2 proteins which harbored *trans*-acting hotspots. (B) Mcr1 protein levels as a function of F6 progeny genotypes at the *PDE2*, *IRA2*, *IRA1*, and *MCR1* loci, as indicated. Boxes show median and upper and lower quartiles; whiskers show 1.5 times the interquartile range. (C) tSNE embeddings highlighting proteins upregulated by the vineyard (blue) and clinical (orange) alleles of *IRA1*, *IRA2*, and *PDE2*, as indicated. (D) Schematic illustrating the principle of the pQTL sign test. (E) Mean fraction of coherent *trans*-pQTLs across all mapped associations (ordinate) as a function of *trans*-pQTL *p* values (abscissa). Actual mapping data is shown in purple; random expectation across all *trans*- pQTLs, regardless of protein target, is shown in grey; *p* values by binomial test. See also Figure S3.

### Directional selection drives proteomic divergence

Many pQTL mutations occur at high frequencies in natural yeast populations^20^ [**Fig. S3B**]. Strains bearing the *IRA1*^RM^*/IRA2*^RM^ (vineyard) allele combination, for instance, were isolated from strikingly similar ecological niches, including grape must, soil below a rotten apple, Uruguayan wine, Tokay grapes, and orange juice concentrate^20^. To understand the molecular consequences of the Ras/PKA hotspot variants across these backgrounds, we examined their proteomic effects^32^. Our atlas accurately forecasted the effects of the RM and YJM *IRA1* and *IRA2* genotypes across the wild isolates: the differences in protein levels between strains with *IRA1*^RM^*/IRA2*^RM^ and *IRA1*^YJM^*/IRA2*^YJM^ genotypes agreed well with mapping predictions [**Fig. S3C**]. Thus, just as for the *cis*-acting variants above, *trans* regulatory effects identified in the F_6_ segregant panel are highly penetrant across other genetic backgrounds, despite wild strains harboring hundreds of thousands of other variants.

The convergence of the pleiotropic hotspots and their evident effects across divergent yeast isolates suggested that selection might have driven polygenic adaptation *via* these mutations, with one set of niches favoring higher expression of the RM-upregulated module and another the YJM- upregulated module. We formalized this hypothesis in a variation on Orr’s sign test^45^, in which we calculated the fraction of pQTLs impinging on a given protein that acted in the same direction [**Fig. 3D**]. We compared this statistic to the null hypothesis that the extent of coherence (the fraction of pQTL-pQTL pairs acting on a given protein that have the same sign) should be no greater than the average coherence across all variant-protein associations. A significant deviation in the observed extent of coherence suggests that we can reject neutrality and conclude that directional selection acted to shape the concerted action of *trans*-pQTLs.

Strikingly, the effects of pQTLs on protein level were much more coherent than expected by chance (binomial test *p* < 10^-250^ for pQTL-pQTL pairs with *p* values < 10^-10^). The coherence was pronounced across a wide range of pQTL *p* value thresholds [**Fig. 3E**], and the trends we observed were driven by both RM-higher and YJM-higher coherent pQTL-pQTL pairs [**Fig. S3D**]. These data indicate that the RM and YJM parental backgrounds have undergone directional selection on the expression of these proteins, driven by multiple variants controlling the same regulatory modules. The coherence in *trans*-pQTL effects we observed, therefore, is likely adaptive and ecologically relevant.

### Coding variation driving protein abundance in trans

Protein abundance can be controlled either by coding (protein-altering; non-synonymous) or non-coding (regulatory, and also potentially synonymous) mutations either in *cis* or in *trans* [**Fig 4A**]. Both coding and non-coding variants altered protein abundance in *cis*: just under half of the *cis*- acting pQTNs we identified altered protein-coding sequences, and both protein-altering and regulatory variants had similar effect sizes [**Fig 4B**]. On the other hand, protein-altering *trans*- pQTNs exerted much larger effects on their targets [**Fig. 4C**]. The Asn201Ser missense variant in Ira2, for instance, was identified in our map to strongly affect the abundance of Mcr1 (among many other targets) [**Fig. 3B**]. To confirm that this variant was causally responsible, we reconstructed the allele of the clinical strain, by introducing the single, trans-acting nucleotide variant in the vineyard strain background by genome editing. We observed a pronounced decrease in Mcr1 levels (*p* < 0.0002) also in the vineyard background [**Fig. 4E**]. Thus, the homeostatic network of cells may buffer the proteomic effects of regulatory *trans*-pQTNs relative to their protein-coding counterparts.

**Figure 4.**
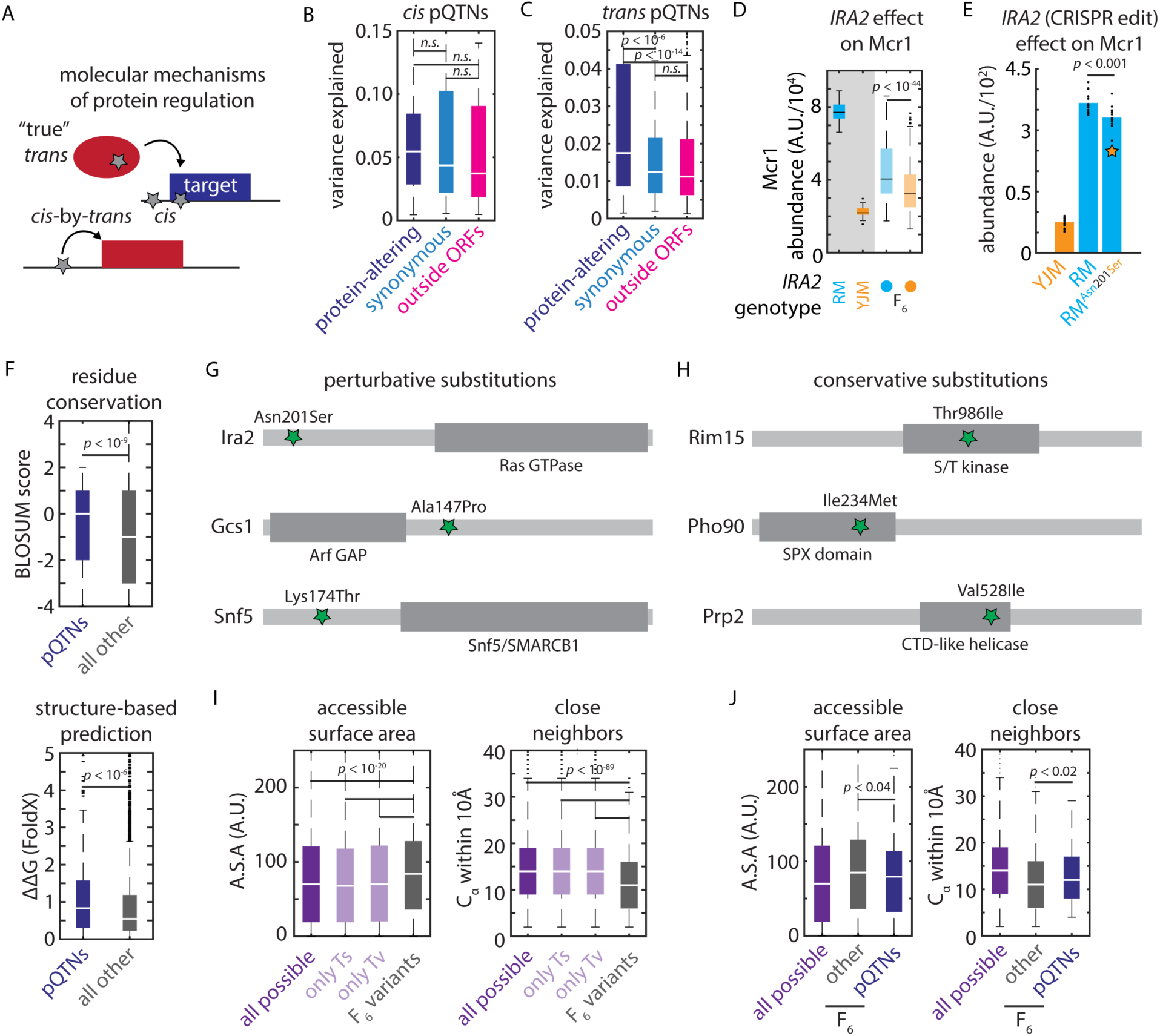
Biochemical constraints revealed by proteomic mapping. (A) Schematic illustrating possible molecular mechanisms of *cis* and *trans* regulation (B) Effect size of protein-altering, synonymous, and regulatory *cis*-pQTNs, as indicated. Boxes show median and upper and lower quartiles; whiskers show 1.5 times the interquartile range. (C) Effect size of protein-altering, synonymous, and regulatory *trans*-pQTNs, as indicated. Boxes show median and upper and lower quartiles; whiskers show 1.5 times the interquartile range. *p* values by two-sided *t* test. (D) Predicted effect from genetic mapping of the *IRA2*^Asn201Ser^ missense variant on Mcr1 levels. *p* value by *F* test. (E) CRISPR reconstruction and mass spectrometry to validate the effect of the *IRA2*^Asn201Ser^ variant on Mcr1 levels. *n* = 15; *p* value by two-sided *t* test. (F) BLOSUM62 (top) and FoldX scores (bottom) for missense *trans*-pQTNs (blue) as compared to all other segregating missense variants (grey). Boxes show median and upper and lower quartiles; whiskers show 1.5 times the interquartile range. *p* values by Mann-Whitney *U* test. (G) Illustrative conservative pQTN substitutions and (H) perturbative pQTN substitutions with functional domains of the mutated proteins indicated. (I) Solvent-accessible surface area and number of C_α_ within 10Å for all possible missense SNPs (purple; also shown are subsets resulting from transitions and transversions) and all missense variants segregating in the F_6_ mapping panel (grey). (J) As in (I) for all possible missense SNPs (purple), missense pQTNs identified in this study (blue), and all other missense variants segregating in the F_6_ mapping panel (grey). *p* values by Mann-Whitney *U* test. See also Figure S4.

The strength of these effects led us to speculate that coding *trans* pQTNs–which perturb the protein products of the genes in which they arise–might help us to understand the biochemical features of missense variants that impact function. We first used a classic metric (BLOSUM62^46^) to assess the conservation of missense *trans* pQTNs as compared to all other segregating missense variants. To our surprise, missense pQTNs were more conservative (in terms of BLOSUM62 score) than the control variants (*p* < l0^-9^) [**Fig. 4F**], suggesting that knowledge of the reference and alternate amino acid residues was insufficient to predict functional outcomes. With this in mind, we used the FoldX variant effect prediction algorithm–which incorporates protein structures–to score the pQTNs and the set of control missense variants^47^. This analysis indicated that *trans* pQTNs were indeed more disruptive to protein stability than other segregating missense mutations (median ΔΔG ∼ 0.83 vs. 0.54 kcal/mol; *p* < 10^-6^) [**Fig. 4F**].

The discrepancy between the BLOSUM62 and FoldX predictions suggested that local context within a protein was important. Consistent with this idea, amongst the pleiotropic *trans* hotspots we identified, perturbative missense *trans*-pQTNs often occurred outside of the core functional domains of the encoded protein: Ira2^Asn201Ser^ (356 targets) lay outside of the Rho GTPase domain; Gcs1^Ala147Pro^ (50 targets) outside of the ArfGAP catalytic domain; and Snf5^Lys174Thr^ (37 targets) in a disordered region outside of the conserved SNF5/SMARCB1 domain [**Fig. 4G**]. The opposite was true for conservative substitutions: Rim15^Thr986Ile^ (36 targets) lay in the kinase domain; Pho90^Ile234Met^ (31 targets) in the SPX domain; and Prp2^Val528Ile^ (24 targets) in the helicase domain [**Fig. 4H**]. We also noted that Ura5^Gly73Val^, which controlled 79 targets, lay in the core phosphoribosyltransferase domain of the enzyme – this may account for its strong and widespread effects.

Generalizing this idea, we hypothesized that two parameters might capture key aspects of the structural context: 1) a residue’s solvent-accessible surface area and 2) the number of other alpha-carbon atoms within 10Å (a proxy for the local complexity of the protein fold). Together, we expected these metrics to capture the proximity of a residue to a protein’s core folded and functional domains. Exploiting the availability of AlphaFold2-predicted backbone structures^48^, we calculated these statistics for every residue in the yeast proteome. Reasoning that missense variants that fixed in wild strains might themselves represent a conservative subset of the possible mutational spectrum, we first compared all segregating missense variants in our cross to all possible missense SNPs that could arise in the proteome (see **Methods**). Indeed, the mutations present in the F_6_ progeny used in our experiments were both more solvent-exposed and occurred in less-complex regions of the protein fold (*p* < 10^-20^; *p* < 10^-90^; respectively) [**Fig. 4I**; **Fig. S4A**]. The same was true when considering only transitions or only transversions, suggesting that this finding was independent of biases in the origin of the natural mutations. Nevertheless, amongst these fixed mutations, both structural metrics distinguished missense pQTNs from all other segregating missense variants: pQTNs were more buried and occurred in more complex regions of the fold relative to other segregating variation (*p* < 0.04; *p* < 0.02; respectively) [**Fig. 4J**]. Collectively, these data illustrate how nucleotide-resolution genotype-to-molecule maps can reveal biochemical mechanisms changing protein abundance and, in turn, explain the prevalence of natural genetic variants.

### Covariation of protein abundances reveals foundational proteome architecture

Precise deletion and knockdown experiments yield rich information on the molecular and functional connectivity of gene products^30,49,50^, but remain challenging in non-model organisms and for essential genes. In large proteomic datasets, protein covariation analysis is a powerful alternative strategy to learn about protein function, and is particularly effective for essential proteins, which are enriched for high abundance and low variability^51^. We first calculated the correlation in protein abundance across our mapping cohort for all pairs of observed proteins, noting many covariation signals that reflected known metabolic functionality. For instance, levels of Hxk2, the glycolytic hexokinase that predominates during growth on glucose, were strongly anticorrelated with its paralog Hxk1 and the hexokinase Glk1 [**Fig. S5A**]. Both Hxk1 and Glk1 are directly repressed by nuclear localization of Hxk2 under low glucose concentrations^52^. Hxk1 and Glk1 levels were themselves tightly correlated, as was Emi2, a paralog of Glk1 with hexokinase activity^53^. Indeed, these relationships were reflective of the broad tradeoff between fermentative and respiratory gene expression programs: glycolytic and citric acid cycle enzymes [**Fig. 5A**] were coherently controlled by the *IRA2* alleles described above [**Fig. 5B**]. These regulons formed pronounced covarying clusters [**Fig. 5C**]; notably, this covariation structure was much more evident amongst the F_6_ progeny than in biological replicates of the parents alone [**Fig. 5D**].

**Figure 5.**
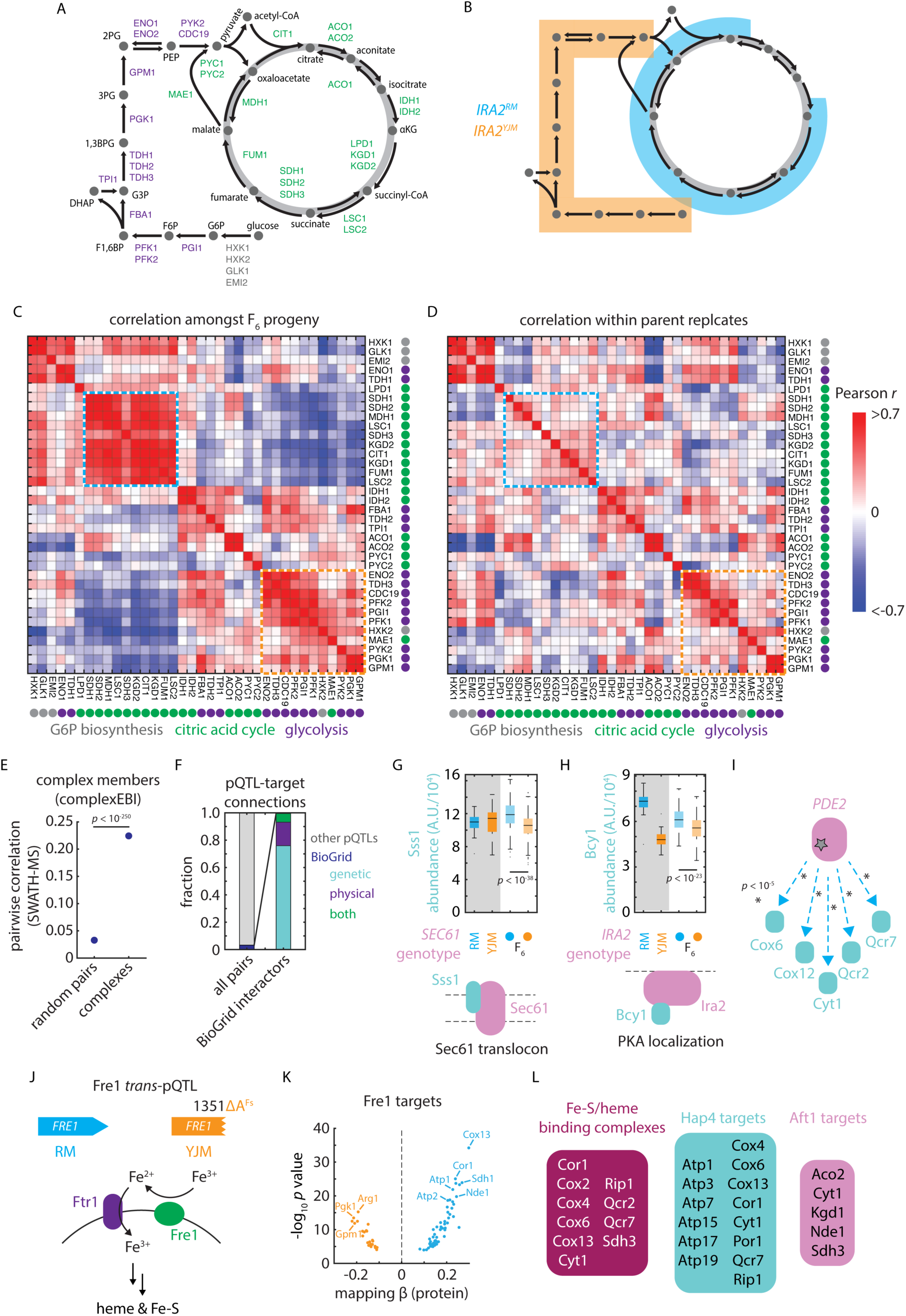
pQTLs reveal molecular and functional connectivity. (A) Schematic of metabolites and enzymes of glycolysis (purple) and citric acid cycle (green). (B) As in (A), with metabolites highlighted in blue and orange if an enzyme catalyzing a reaction involving that metabolite is regulated by *IRA2^RM^*or *IRA2^YJM^* alleles, respectively. (C) Heatmap of pairwise SWATH-MS abundance correlations amongst enzymes shown in (A). Highlighted in blue and orange are blocks of coregulated enzymes regulated by the *IRA2^RM^* or *IRA2^YJM^*alleles, respectively. (D) As in (C), but for correlations within replicate measurements of parental isolates. (E) Pairwise SWATH-MS abundance correlations between complex members as compared to all possible pairs of measured proteins. *p* value by Mann-Whitney *U* test. Dots indicate mean and bars standard error. (F) Cumulative frequencies of pQTL-target connections reflecting (left) BioGRID interactions (blue) and all other pQTL-target pairs (grey) and (right), amongst BioGRID interactions, those annotated as genetic (blue), physical (purple) or both genetic and physical (green). (G) Sss1 abundance in vineyard and clinical parents and in F_6_ progeny with *SEC61* genotypes as indicated. (H) Bcy1 abundance in vineyard and clinical parents and in F_6_ progeny with *IRA2* genotypes as indicated. (I) Schematic of pQTL-target connections between *PDE2* and various targets upregulated by vineyard allele, as indicated. *p* values by *F* test. (J) Schematic of the role of Fre1 in iron reduction and uptake at the plasma membrane ^67^. (K) Volcano plot illustrating predicted effects on abundance from genetic mapping (abscissa) and forward selection *F* test *p* value (ordinate) for the *FRE1 trans-* pQTL. (L) Downstream *FRE1* pQTL targets that bind iron or heme or that are targets of Hap4 or Aft1, as indicated. See also Figure S5.

We then asked whether covariation in these closely related F_6_ progeny was representative of covariation across natural and synthetic genetic diversity in *S. cerevisiae*. We compared the correlations in protein abundance in our dataset to those in a species-wide survey^54^, as well as the correlations observed within the proteomes of ∼ 5,000 viable gene deletion strains^30^. The architecture of covariation was conserved, with protein covariation coefficients correlating well between these independent experiments (Pearson’s *r* = 0.56 for F_6_ haploids vs. 1,002 Yeast Genomes; 0.52 for F_6_ haploids vs. precise deletions) [**Fig. S5B**]. Thus, the modest genetic divergence harbored by our mapping panel drives proteome diversity that is representative of a much broader range of genetic variation.

### Systems biology of variant-protein associations

To probe the physical and genetic connections embedded in these data, we first assessed whether members of the same macromolecular complex^55^ co-varied in their abundance. Indeed, the mean Pearson correlation between complex members was 0.224, as compared to 0.038 for all protein-protein pairs (*p* < 10^-195^ by Mann-Whitney *U* test) [**Fig. 5E**]. These data were sufficient to resolve the fine details of complexes and metabolic pathways: we found, for instance, that the F_1_ core structural subunits (particularly the alpha (Atp1), beta (Atp2), gamma (Atp3), and a component of the stator (Atp4) of the mitochondrial ATP synthase) were highly correlated [**Fig. S5C**]. Similarly, the levels of enzymes with functional overlaps or that physically associate (*e.g.* Idh1/Idh2, Kgd1/Kgd2) covaried tightly [**Fig. S5D**]. Abundance correlations were also reflective of other measures of connectivity. The STRING database co-expression metric, which aggregates mRNA and protein data^56^, was significantly correlated with protein covariation in our measurements (*p* < 10^-250^) [**Fig. S5E**]. So too was the genetic interaction similarity score from The Cell Map (*p* < 10^-191^)^57^ [**Fig. S5F**], which measures functional relatedness based on genetic epistasis analysis.

Protein covariation can be caused by physical interactions between proteins. We thus speculated that some of the architecture of our mutation-to-protein atlas could be mechanistically explained by interactions between complex subunits (from ComplexEBI^58^) and genetic or protein-protein interactions (obtained from BioGRID^59^) [**Fig. 5F**]. Only one *trans*-pQTL connected two members of the same complex: Sss1 and Sec61 participate in the conserved Sec61/SecYEG translocon complex and, notably, Sss1 plays a key role in the stability of the Sec61 protein [**Fig. 5G**]^60^. A further 204 (∼ 3.2%) *trans*-pQTL-target pairs connected protein-protein interactors [**Fig. 5F**]. Of these, 155 were genetic interactors, 35 physical, and 14 both genetic and physical. A variant in *IRA2*, for instance, controlled the abundance of the PKA regulatory subunit Bcy1; these proteins physically interact as part of the Ras/PKA signaling complex [**Fig. 5H**]^61^. Similarly, a variant at *PDE2* controlled an array of its genetic interactors, including Cox6, Cox12, Cyt1, Qcr2, and Qcr7, all of which are involved in respiration–a process tightly linked to cAMP signaling mediated by Pde2 [**Fig. 5I**]^39^. Thus, *trans*-pQTL relationships reflect known physical and functional associations between proteins, while also describing a rich regulatory network not captured by complementary interaction metrics.

### Functionalizing the proteome reveals cryptic regulatory activity

A surprising example of these noncanonical regulatory networks arose at *FRE1*, a gene encoding a ferric reductase important in iron and copper uptake and metabolism [**Fig. 5J**]^62^. The pleiotropic hotspot, attributable to a frameshift in *FRE1* in the clinical (YJM) background, controlled the levels of 79 proteins (56 upregulated by the vineyard allele and 23 by the clinical allele) [**Fig. 5K**]. Only 2 of the regulated genes exhibited genetic interactions with *FRE1* in BioGRID, and none were physical interactors. Strikingly, however, many of the targets and their associated complexes depended on heme or iron-sulfur clusters for their activity (*e.g.*, Cor1, Cox2/4/6/13, Cyt1, Qcr2/7, Rip1, Sdh3) or were otherwise involved in respiration (*e.g.*, Atp1/2/3/5/7/15/17/19, Cit1, Fum1, Kgd1/2, Mdh1, Sdh1/3) [**Fig. 5L**]. Indeed, iron metabolism and mitochondrial function are intimately linked^63^.

The downregulated set of proteins was also highly enriched (*p* < 10^-19^)^36^ for targets of the *H*eme *A*ctivator *P*roteins (Hap) 2/3/4/5 transcription factor complex, which respond to intracellular heme levels^64,65^. Conversely, the set of proteins upregulated in the *FRE1* loss-of-function background were enriched for targets of Nhp6 (*p* < 10^-4^), which acts with Aft1 (*A*ctivator of *F*errous *T*ransport) in the upregulation of iron transport^66^. Thus, impaired heme and iron-sulfur cluster synthesis–due to loss of Fre1 activity–led to widespread downregulation of enzyme components that depend on iron to function and an upregulation of compensatory transport machinery. The ubiquity of these noncanonical hotspots in our atlas suggests that connecting mutations to molecules can reveal previously unappreciated regulatory relationships–indeed, some may be mediated directly by cofactors or metabolites.

### Prioritizing causal variants at drug-resistance loci

Variants that impact molecular phenotypes are often thought more likely to underlie organismal traits. A promising application of mutation-to-molecule maps is therefore to prioritize causal variants at poorly resolved loci that are implicated by genotype-to-phenotype mapping (*e.g.*, based on GWAS)^9^. To assess the validity of this heuristic in our real-world dataset, we analyzed a complementary high-resolution genotype-to-phenotype map^34^ across an array of carbon sources, antifungal drugs, mutagens, and toxic metals. Across 12 environments, we mapped 9,321 QTLs and resolved 2,519 QTNs to a single causal variant (FDR ∼ 10%; see **Methods**), explaining a median of 64% of the phenotypic variance at the final experimental time point [**Supplemental Table S7**].

We noted that the RM allele of a regulatory variant (*ERG11^T122014C^*) adjacent to *ERG11* was predicted to upregulate the associated protein Erg11, the mechanistic target of the azole antifungals in *S. cerevisiae*^68^, and to reduce sensitivity to azole treatment. Yet our phenotypic mapping also implicated a missense variant, Erg11^Lys433Asn^, as potentially important for fluconazole sensitivity–albeit without resolving the mutation as a phenotypic QTN [**Fig. 6A**]. Upon reconstruction of these mutations in the sensitive background by genome editing, mass spectrometry confirmed that the upstream regulatory variant controlled protein level, as predicted [**Fig. 6B**]. The neighboring missense variant, as expected from our mutation-to-protein map, did not impact abundance. Both of the variants, however, reduced azole sensitivity in additive fashion (*p* < 0.05) [**Fig. 6C**]; thus, the combination of proteomic and phenotypic mapping revealed two variants at this locus that contribute equally to drug susceptibility. This example and others^22,35,69^ emphasize that the architecture of even a single causal locus can be complex, and that non-coding variation cannot be neglected when identifying and predicting drug resistance^70^.

**Figure 6.**
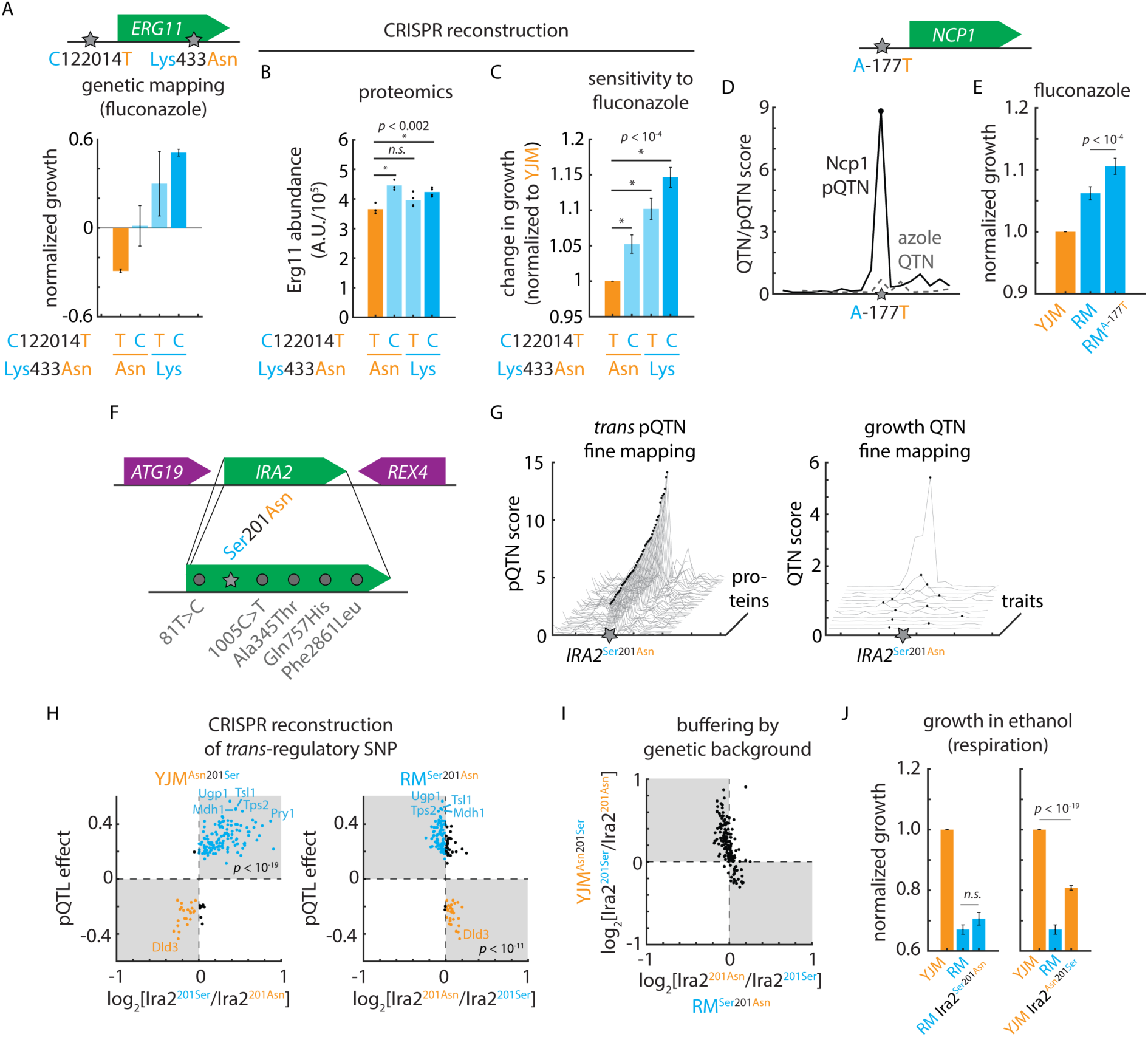
Cryptic fitness effects embedded in the mutation-to-protein map. (A) Genetic mapping of the phenotypic effects of *ERG11^T1220124C^* and Erg11^Asn433Lys^ in fluconazole. Shown is normalized growth of F_6_ progeny with genotypes as indicated. (B) Mass spectrometry of Erg11 protein levels in clinical (YJM) wild-type and CRISPR-edited YJM *ERG11^T1220124C^*, YJM Erg11^Asn433Lys^, and YJM *ERG11^T1220124C^* Erg11^Asn433Lys^ mutant strains. *n* = 4; *p* values by Student’s *t* test. (C) Growth of clinical (YJM) wild-type and CRISPR-edited YJM *ERG11^T1220124C^*, YJM Erg11^Asn433Lys^, and YJM *ERG11^T1220124C^* Erg11^Asn433Lys^ mutant strains in fluconazole. *n* = 96; *p* values by Student’s *t* test. (D) Fine-mapping of Ncp1 *cis*-pQTN as compared to fine-mapping of the azole-sensitivity QTL in the vicinity of *NCP1*. (E) Growth of clinical (YJM), vineyard (RM), and CRISPR-edited RM *NCP1^A-177T^*mutant strains in fluconazole. *n* = 96; *p* value by Student’s *t* test. (F) Diagram of *IRA2* locus and segregating *IRA2* mutations. (G) pQTN fine-mapping scores for the top 50 *IRA2*-target associations (left) and QTN fine-mapping scores for *IRA2* growth QTL associations. (H) Predicted *IRA2* pQTN effects from genetic mapping (this study; ordinate) as compared to measured effects of (left) CRISPR-edited YJM Ira2^Asn210Ser^ and (right) RM Ira2^Ser201Asn^ mutants. Mass spectrometry estimated abundances normalized to wild type in each case. (I) Measured effects of CRISPR-edited YJM Ira2^Asn210Ser^ (ordinate) and RM Ira2^Ser201Asn^ (abscissa) mutants. (J) Growth of clinical (YJM), vineyard (RM), and CRISPR-edited RM Ira2^Ser201Asn^ mutant (left) and YJM Ira2^Asn210Ser^ mutant (right) in ethanol. *n* = 96; *p* values by Student’s *t* test. See also Figure S6.

We examined our growth mapping data for other examples to support the notion that unresolved genotype-to-phenotype associations could be resolved by proteogenomic mapping. One striking example was the non-coding *cis*-pQTN *NCP1*^A-177T^. Ncp1 associates with the ergosterol biosynthetic enzyme Erg11, and phenotypic mapping suggested that a causal locus for fluconazole sensitivity was present, but we failed to implicate a single QTN [**Fig. 6D**]. Yet when we reconstructed the putative causal variant and subjected the gene-edited strain to azole treatment, the higher-expressing *NCP1*^-177T^ allele indeed exhibited decreased azole sensitivity (*p* < 10^-4^) [**Fig. 6E**]. The *NCP1* mutation did not impact growth in the absence of drug [**Fig. S6A**], nor significantly increase the levels of Erg11 [**Fig. S6B**], indicating that the effect on azole sensitivity was likely directly related to Ncp1 abundance. These case studies illustrate how proteogenomic mapping can inform detailed hypotheses regarding the function of natural variants.

### Molecular mapping pinpoints a hidden causal variant

*Trans*-regulatory mutations are often thought to have widespread effects on phenotype due to changes in the expression of many downstream target proteins^16^. Considering the large number of proteins–more than 300–regulated by the *IRA2* hotspot, we anticipated a strong phenotypic effect. To our surprise, however, QTN mapping revealed few variant-phenotype associations at *IRA2*, even though dozens of pQTNs were unambiguously identified [**Fig. 6FGH**; **Fig. S6C**]. To understand this discrepancy, we first confirmed that the numerous variant-protein associations at the *trans*-pQTL hotspot reflected a change in Ira2 and not a linked mutation in a neighboring gene. Comparing our mapping results to orthogonal proteomic characterization of an *IRA2* deletion allele^30^ strongly suggested that the hotspot was attributable to loss of Ira2 function: the proteomic consequences of the YJM allele of *IRA2* were highly concordant with those of the deletion (*r* = - 0.81; *p* < 10^-80^; *i.e.*, the RM allele is hyperactive) [**Fig. S6D**]. Much weaker correlations were observed between our mapping data and the proteomic effects of deleting the neighboring *ATG19* and *REX4* genes (*r* = −0.08 and *r* = 0.30, respectively).

We next tested whether the Ira2^Asn201Ser^ mutation alone, and not one of the several other mutations segregating at *IRA2*, was responsible for the predicted regulatory effects. Reconstructing the putative causal variant had widespread effects on protein abundance that agreed very well with our mapping results: nearly all the proteomic effects in the clinical (YJM) background (94%; *p* < 10^-19^) and the majority in the vineyard (RM) background (78%; *p* < 10^-11^) agreed with the mapping prediction [**Fig. 6H**]. Thus, the Ira2^Asn201Ser^ mutation is a true pleiotropic *trans*-pQTN.

Although highly sensitive, our phenotypic and pQTL mapping approaches (like most, with a handful of exceptions, *e.g.*^71^) assume a linear model in which the effects of mutations combine additively. We therefore entertained the possibility that while the regulatory effects of the Ira2^Asn201Ser^ mutation were as predicted, its effects were modified by nonlinearities not captured by our linear model (*e.g.*, those arising due to genetic background effects). Indeed, the quantitative consequences of the *IRA2* variant were much more pronounced in the YJM background than in its RM counterpart, despite widespread directional concordance [**Fig. 6I**]. This suggested that a genetic background effect might be at play.

With this in mind, and considering that Ras/PKA signaling is central to nutrient sensing, we measured the growth of the genome-edited strains bearing the *trans*-regulatory mutation on various carbon sources. Strikingly, we found that the Ira2^Asn201Ser^ mutation had fitness effects that were both strain- and condition-specific: the vineyard allele was highly deleterious in the clinical background when cells were grown on non-fermentable carbon sources, whereas the clinical variant had a minimal fitness effect when reintroduced into the vineyard background [**Fig. 6J**]. Conversely, the clinical mutation modestly impacted fermentative growth in the vineyard background, while the vineyard mutation had no significant effect under such conditions in the clinical parent [**Fig. S6E**]. The asymmetric phenotypic effects of the Ira2^Asn201Ser^ polymorphism were likely obscured in statistical mapping due to the segregation of suppressing alleles responsible for the strong background effect. Thus, molecular mapping can unmask nonlinearities that otherwise disguise the fitness effects of even highly pleiotropic regulatory hotspots, and forecast their impact under the conditions where these effects emerge.

### Forecasting variant effects across environments from molecular phenotypes

The proteomic measurements that determined our mutation-to-protein map were made only in the absence of stress: the causal *cis*-regulatory variant at *ERG11* for instance [**Fig. 6A**], was identified as a pQTN in media without azoles, but was a potent regulator of azole sensitivity. Likewise, the widespread proteomic impact of the *IRA2* hotspot mutation was readily apparent in minimal glucose, even though its fitness consequences emerged more strongly in respiratory conditions. Moreover, stress-response QTLs were highly condition-specific: we saw little decline in the identification of unique QTLs as we considered additional environments [**Fig. 7A**]. In concordance with an omnigenic model of complex heritability in which many genes contribute to a phenotype^72^, stress-response traits were more genetically complex than protein levels and stress-response QTLs exhibited smaller median effect sizes than pQTLs (*p* < 10^-16^) [**Fig. 7B**]. Consistent with their larger effects, and in contrast to phenotypic QTLs, rarefaction analysis indicated that we captured a comprehensive pQTL atlas [**Fig. 1K**]. Together, these properties suggested that the genotype-to-protein map was well-powered to dissect the molecular mechanisms underlying emergent stress-response QTLs.

**Figure 7.**
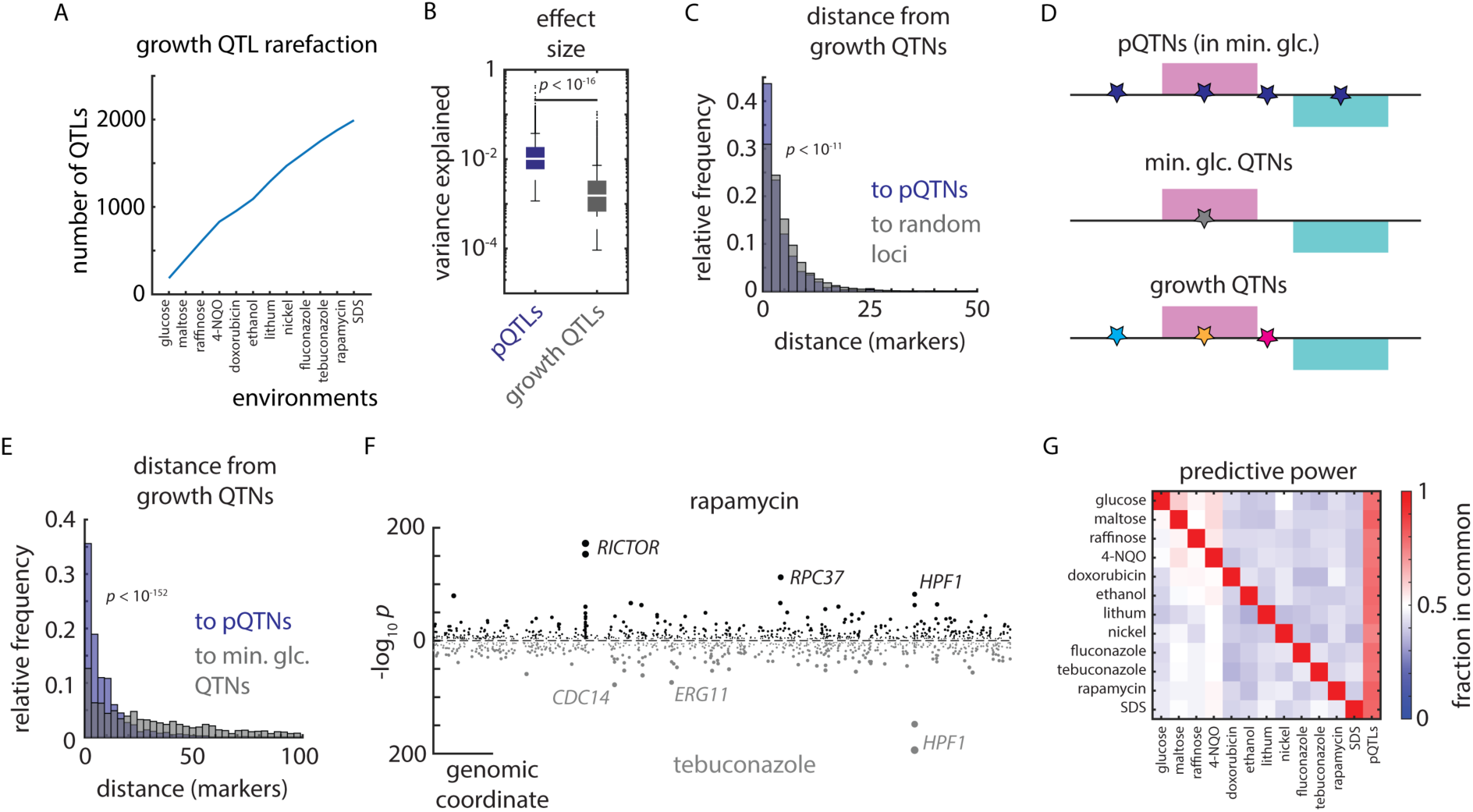
Proteomes identify causal variants underlying quantitative traits. (A) Rarefaction plot of unique growth QTLs discovered as a function of additional environments mapped, as indicated. (B) Effect size (variance explained) of pQTLs (blue) and growth QTLs (grey). *p* value by Mann-Whitney *U* test. (C) Relative frequency histogram of the distance from al phenotypic QTNs to (blue) the nearest pQTN and (grey) randomly selected sets of markers of the same size. *p* value by Kolmogorov–Smirnov test between real and permuted data. (D) Schematic of pQTNs (blue), growth QTNs in minimal glucose medium (no stress; grey), and stress-responsive growth QTNs (various colors). (E) As in (C), but illustrating the distance from stress-responsive growth QTNs to (blue) the nearest pQTN and (grey) growth QTNs discovered in minimal glucose (no stress). *p* value by Kolmogorov–Smirnov test. (F) Example Miami plot of QTLs identified for growth in rapamycin (top) and tebuconazole (bottom). (G) Heatmap of the relative fraction of QTLs in common between environments (ordinate) and environments and pQTLs (abscissa), as indicated. See also Figure S7.

Although effects on phenotype arise through diverse mechanisms, and only a subset act *via* changes in protein level, our mutation-to-protein map overall contained rich information on causality: pQTNs were much closer to stress-response QTNs than expected by chance (*p* < 10^-11^) [**Fig. 7C**]. Moreover, the effect sizes of pQTNs and stress-response QTNs were correlated (*r* = 0.29; *p* < 10^-16^) [**Fig. S7A**]. We therefore hypothesized that growth phenotypes reflect the effects of underlying mutations controlling proteins with distinct phenotypic consequences [**Fig. 7D**]. Indeed, across all the environments we surveyed, pQTNs discovered in minimal glucose medium were much more predictive of causality under stress (*p* < 10^-152^) than QTNs from the minimal glucose condition [**Fig. 7E**].

Rapamycin and tebuconazole resistance traits, for instance, shared few large-effect QTLs in common (*e.g. HPF1*) and were predominantly driven by distinct loci (*e.g.* at *RICTOR/AVO3* and *ERG11*, respectively): of the 663 causal loci identified in rapamycin, only 268 (40%) coincided with one of the 635 causal loci underlying tebuconazole resistance [**Fig. 7F**]. Strikingly, on the other hand, the genetic architecture of protein levels had much greater overlap, with 489 tebuconazole-resistance loci (77%) and 534 rapamycin-resistance loci (81%) coinciding with a pQTL. This was true in general: across the diverse environments we tested, an average of 78% of stress-resistance QTLs colocalized with a pQTL, in contrast to an average of only 48% of QTLs coinciding between stress conditions [**Fig. 7G**]. This suggested that the molecular effects of genetic diversity that pre-existed in unstressed cells emerged into distinct cellular phenotypes under stress. Molecular mapping therefore holds powerful promise in forecasting the functional consequences of mutations, even if the maps are charted before pathologies emerge.

## Discussion

Most mutations, and even many genes, remain of unknown cellular function. A promising bridge from genotype to phenotype is to map the effects of natural variants on protein levels, because proteins perform an array of critical cellular functions that link the DNA blueprint to physiology. Despite this promise, causally linking individual mutations to their proteomic consequences and phenotypic effects remains a challenge. This is in part because most genetic mapping approaches yield (at best) gene-level resolution, and also because mutations can alter protein function in various ways; for example, many proteins function in protein complexes or larger molecular pathways. Moreover, associating proteins with phenotypes alone often cannot disentangle whether changes in protein levels are truly causal. Here, building on ‘super-resolution’ phenotype mapping using a large segregant panel from two closely related yeast parents^10,13^, we combined this approach with quantification-precise high-throughput proteomics to link genetic to proteomic diversity.

Although we do not quantify all proteins, we capture a large fraction on a molar basis. Quantifying additional marginal proteins would not change the overall regulatory picture we charted, as indicated by rarefaction analysis (although more *cis*-acting loci would likely be identified). Notably, because essential proteins are enriched in the high-abundance protein fraction well-detected by mass spectrometry^30^, our map captures essential proteins particularly well, and thus complements forward and reverse genetic screens. Moreover, because our cross recombines naturally occurring genetic variation, our study complements deep mutational scans that contain many variants not found in nature.

Despite their small genetic differences, the two parental isolates harbor highly diverged, functionally coherent proteomes. While the boundaries set by the parents largely define the proteomes of the offspring, the offspring exhibited substantial proteomic diversification, as well. Exploiting the segregation of the underlying genetic diversity in the F_6_ progeny, we captured genetic control for most proteins in our atlas, with a surprisingly high number of variants impacting protein levels. Thus, fixed natural mutations were often far from neutral: even the variation between two closely related strains proved to be a rich vein of diversity in the proteome.

Notably, proteins that did not differ in abundance between the parents often changed in the offspring. Termed transgression, this property has been reported for mRNA abundance and organismal phenotypes^10,11,73^, but has thus far received limited attention in the proteome. Similar effects also likely underlie the phenotypic transgression commonly observed in agricultural genetics^73^. Further, for several proteins, their abundance in most of the offspring closely resembled one parent rather than the other. Interestingly, the deviating parent often represented an extreme relative to other wild isolates, while the typical offspring more closely resembled the average of natural strains across the species. Both phenomena are explained in our data by multiple loci that aggregate in controlling the abundance of a protein. Due to different variants driving abundance in opposing directions, the extremes become less likely–but not inaccessible–compared to typical protein levels. Natural genetic diversity is thus amplified in the proteome through meiosis; this emergent proteomic diversity could be a potent source of variation allowing some offspring to rapidly adapt to new environments.

Our dataset demonstrates the added value of proteomics in interpreting genetic variation. We achieved explanatory power previously reported only for mutation-to-mRNA maps^12^, but with the critical addition of very high resolution–often implicating single causal nucleotides. This highlighted the complementarity of eQTL and pQTL approaches: many *cis*-pQTL effects are detected only at the proteome level, with no evidence of mRNA allelic imbalance for the associated mRNAs. These associations likely stem from protein properties that are not represented at the mRNA level, such translation efficiency, protein stability, and turnover. Conversely, we observed widespread signatures of *trans* mRNA regulation in our pQTL map (for example, downstream of the Ras/PKA pathway). The remaining missing heritability in our and prior studies likely arises from a large number of small-effect variants, some epistatic contributions, and epigenetic influences, such as prions, that we have not yet tracked in the meiotic progeny. Previous difficulties in identifying signatures of mRNA-level effects in proteomes likely arose primarily from comparatively limited statistical power to identify and colocalize *trans-*eQTLs and -pQTLs^33,74^. Of note, much of *trans* regulation arose from proteins not usually thought of as regulatory, and illustrates the profound self-regulatory structure of metabolism. Indeed, *trans*-regulatory variants were often found in metabolic enzymes and transporters.

The sheer number of well-resolved pQTLs we identified, and our choice to study the progeny of two wild isolates, rather than one wild and one domesticated strain, granted excellent statistical power to assess natural selection on protein levels. Indeed, a sign test on pQTLs revealed that directional selection had acted to reshape the proteome to fit the niches inhabited by each parent, despite their relatively recent evolutionary divergence. This further suggests that the levels of many proteins are relevant to fitness and subject to selection. These and many prior observations^75^ call into question the notion that much of segregating genetic variation is functionally neutral, or nearly so. Rather, natural proteomes likely reflect an intricate interplay between stabilizing selection–as evidenced by transgression–and directional selection–as reflected in the striking proteomic divergence of the parents and the results of the pQTL sign test.

*Cis*-acting effects were balanced between coding and non-coding variants, but coding variation appears to have a privileged role in *trans* regulation of protein levels. This may be because coding variants can alter both protein function and abundance, while non-coding variants are expected to leave the former untouched. Given that few genes are haploinsufficient, whether in yeast or humans, tolerance of small excursions in the amount of a gene product may be a general property. Indeed, in a recent study addressing aneuploid gene dosage in natural strains, we made similar observations and hypothesized that the attenuation of *trans*-acting regulatory variation may arise from buffering of the proteome against gene expression noise^32^. Accounting for the structural context of the *trans* pQTNs in their host proteins revealed molecular signatures that distinguished pQTNs from other segregating variants. Thus, the potency of a coding *trans*-pQTN likely depends on the amino acid substitution it encodes and the function of the protein domain in which the mutation occurs. Given that less stable proteins are more quickly degraded, we speculate that many pQTNs altered protein abundance by reducing stability. Mapping other molecular layers (*e.g.* metabolite levels) may help to disentangle effects on protein stability *versus* catalytic activity, as may considering the position and role of proteins in the metabolic network, as we have shown elsewhere on longer timescales of adaptation^76^. Another intriguing question for future study is which mutations represent a simple modulation (modest gain or loss) of existing activity *versus* an incipient neofunctionalization or gain of new targets.

Abundance covariation amongst the progeny revealed a rich map of functional associations–much more so than considering covariation only in the parents. The pQTL map revealed functional connections not captured by prior interaction networks, even in yeast where these resources are most complete. In a few cases (< 4%), pQTLs reflected known physical or genetic interactions between the proteins, but to a much larger extent our molecular map reflected physiological interactions not captured by these metrics. These included global metabolic traits, such as a cryptic causal variant in *IRA2*, common in natural strains, which affected the respiration/fermentation balance via the Ras/PKA pathway; functional metabolic traits, such as the iron- and iron-sulfur co-dependency of the respiratory chain; and local metabolic traits such as the anti-correlation of hexokinases. Additional functional relationships can likely be identified by extending high-resolution mapping to post-translational modifications^18^ and protein-protein interactions^77^, which are governed by some overlapping and some distinct processes relative to protein abundance pQTLs. Also of interest is to dissect how many of the surprising pQTL hotspots we identified (*e.g.,* at *FRE1*) are mediated by mRNA levels, or whether they are in part due to direct cofactor binding and posttranslational protein destabilization invisible in the transcriptome. Indeed, cofactors are highly prevalent for several important enzyme classes, such as oxidoreductases (80% having a cofactor) and transferases (36%), highlighting the far-reaching potential of this mechanism^78^. Moreover, there is emerging evidence of many other metabolite-protein interactions that are only beginning to be characterized^79^.

High-resolution molecular mapping also proved valuable in identifying cryptic causal variants hidden in plain sight, such as the epistatic variant we identified in *IRA2*. Indeed, the low SNP density in our mapping panel allowed us to readily pinpoint the function of this mutation, unlike in other genetic backgrounds in which the variant exhibits strong epistasis even within the *IRA2* gene^80^. Proteomics suggests that the phenotypic masking we observed arises in part from buffering of the impact of the mutation across all of its targets. This phenomenon likely arises from multiple suppressor mutations throughout the genome, as in the case of a single segregating suppressing allele we would likely observe a residual phenotypic mapping signal [**Fig. S7B**]. Complex cryptic effects like these are particularly pernicious: they do not manifest as “missing heritability”^81^ but rather as “hidden causality,” because they are suppressed in most progeny. An intriguing area for future investigation is the metabolic basis of this pronounced epistatic effect, and we speculate that genotype-to-protein maps may show the way to many cryptic genetic variants.

Finally, we show that much of the adaptive potential of natural variation under stress can be forecast from molecular genetic mapping: pQTNs that were initially phenotypically buffered were highly predictive of fitness effects in new environments. Thus, proteome diversity may explain emerging phenotypic differences across environments, and may be a mechanistic explanation for the difficulty in predicting phenotype across conditions using genomic data alone. This in turn suggests that molecular maps can highlight variants that are likely to emerge to cause disease even if mutation-to-molecule relationships are mapped before pathologies develop (although such conclusions will likely require the integration of other data, for instance, on which genes are causally related to a pathology). Moreover, data from a single tissue or, as with serum, from a pool of proteins from multiple tissues, likely holds molecular regulatory information to support inferences in other tissues affected by a disease.

## Acknowledgments

We thank Fatma Amari, Kathrin Textoris-Taube, Andrea Lehmann, Christiane Kilian, Daniela Ludwig (all Charite - Universitätsmedizin Berlin) for technical assistance with proteomics sample preparation and measurements. We thank the Jarosz and Ralser Labs for helpful discussions.

This work was supported by the NIH (DP2-GM119140, RF1-AG057334, R01-AG06341801, and R01-HG012366 to D.F.J.; F32-GM125162 to C.M.J.), the National Science Foundation (NSF- MCB116762 to D.F.J.), a Searle Scholar Award (14-SSP-210 to D.F.J.), a Kimmel Scholar Award (SKF-15-154 to D.F.J.), a Vallee Scholar Award (to D.F.J.) and a Discovery Innovation Award from Stanford University (to D.F.J.), Swiss National Science Foundation Postdoc Mobility fellowship 191052 (to J.H.), European Research Council (ERC) under grant agreement ERC-SyG- 2020 951475 (to M.R.), and the Ministry of Education and Research (BMBF), as part of the National Research Node ‘Mass spectrometry in Systems Medicine’ (MSCoresys), under grant agreement 031L0220 (to M.R.). D.F.J. is also a Science and Engineering Fellow of the David and Lucile Packard Foundation. Some of the computing for this project was performed on the Sherlock cluster. We would like to thank Stanford University and the Stanford Research Computing Center for providing computational resources and support that contributed to these research results.

## Author Contributions

Conceptualization, C.M.J., J.H., D.F.J., M.R.; Methodology, C.M.J., J.H., M.M.; Software, C.M.J; Formal Analysis, C.M.J., J.H., P.T.; Investigation, C.M.J., J.H.; Writing – Original Draft, C.M.J., J.H.; Writing – Review & Editing, C.M.J., J.H., D.F.J., M.R.; Visualization, C.M.J., J.H.; Supervision, D.F.J., M.R.; Funding Acquisition, D.F.J., M.R., J.H.

## Supplemental Information

Document S1: Figures S1-S7; Table S6

Table S1: Strain layout for proteomics

Table S2: Protein abundance estimates.

Table S3: Proteins differentially expressed in parental strains

Table S4: pQTL mapping results

Table S5: Allele-specific expression analysis summary

Table S7: Phenotypic mapping results

## STAR Methods

### Resource availability

#### Lead contact

Requests for resources and reagents should be directed to and will be fulfilled by the lead contact, Prof. Daniel F. Jarosz.

#### Materials availability

All strains and plasmids used in this study are available upon request to. The F_6_ haploid mapping panel is also available from NCYC.

#### Data and code availability

Mass spectrometry datasets will be publicly available at the proteomics identification database (PRIDE) upon publication.

All custom genetic mapping and protein structure analysis code is available on GitHub (https://github.com/cjakobson/pqtl-mapping; https://github.com/cjakobson/pop-gen-structure).

Analyses and plots for the figures can be reproduced by cloning the *pqtl-mapping* repository, downloading the contents of the *pqtl-mapping-dependencies* folder (https://www.dropbox.com/scl/fo/3xbcbe9ivwz8aahrlk137/APGxHor01S7jnNX3a1Yk3Og?rlkey=yx81ckrtaq8eb5pu80ggprjhs&dl=0), and running *plotting_master_script.m*.

The dependencies will be deposited at Zenodo upon publication.

### Experimental model details

#### Yeast strains

*Saccharomyces cerevisiae* strains for genetic mapping were generated and genotyped previously as described in^22^. Briefly, ∼1,000 F_6_ progeny from a cross between RM11 and YJM975 were arrayed from single-colony isolates and subjected to whole-genome sequencing. To avoid confounding effects of segregating auxotrophic markers in our proteomics experiments, we selected ∼850 strains from the original panel that were auxotrophic only for uracil (leucine auxotrophy also segregates). In addition to these progeny, we included at least three biological replicates of the RM11 (YDJ6649) and YJM975 (YDJ6635) haploid parental isolates in each 96- well plate of our measurement campaign. These haploid strains are auxotrophic only for uracil to match the F_6_ segregant progeny. Also included were representative haploid wild isolates (22 strains in up to n = 6 replicates) from throughout the world, as cataloged in the SGRP collection^27^. The plate layouts and strain identifiers for the proteomics campaign can be found in **Supplemental Table S1**. A table of other yeast strains used in this study can be found in the **Key Resources Table**.

#### Media and culture conditions

Unless otherwise noted, yeast were propagated in minimal glucose medium with uracil (20 g/L glucose; 6.7 g/L yeast nitrogen base; 20 mg/L uracil; 20 g/L agar as needed for solid medium). Samples for growth phenotyping were pre-grown for 24-48 hr at 30°C on minimal glucose agar with uracil on Singer PlusPlates before replica pinning to growth conditions as indicated using a Singer ROTOR.

For proteomics, samples were spotted from 12×96-well cryo stocks to Singer PlusPlates with 40 ml agar minimal medium using a Singer ROTOR and grown for 4 hours at 30°C. Cells were then transferred with the Singer ROTOR to 96-well plates with 200 µl minimal medium, and incubated for 16 hours. Then, 160 µl of each well of this preculture was transferred to 2 ml wells in a 96-deep-well plate with 1440 µl minimal medium and with one 2 mm borosilicate bead per well. Plates were then sealed with a Breathe Easier sealing membrane (Sigma Aldrich) and incubated on 4 shakers (Heidolph Titramax 1000, 750 rpm, 30°C, 8 hours). After incubation 1.4 ml were transferred to fresh 96-deep-well plates, and harvested by centrifugation (5 min, 4000 g). The supernatant was discarded, plates sealed with adhesive aluminum foils, and the pellets stored frozen until further processing (−80°C). Subsequently, in each well 1600 µl sterilized water was added to the ∼200 µl culture remaining in original incubation plates, plates were quickly vortexed, and OD_600_ was determined using a multi-well plate reader (Spark-Stacker, Tecan). For proteomes of reconstructed strains, samples were prepared in a similar fashion, containing strains YDJ6635, YDJ8281, YDJ8436, YDJ8437 (“batch 1”), YDJ6635, YDJ6649, YDJ8524, YDJ8525, YDJ8526 (“batch 2”) and YDJ6635, YDJ6649, YDJ8527, YDJ8528, YDJ8529, YDJ8578 (“batch 3”).

### Method details

#### Proteomics sample preparation

Frozen pellets were thawed on ice. Segregant samples were processed in 3 batches with 4×96-well-plates each, whereas reconstruction strains were prepared in 96-well plates in their respective sampling batches. To each well/plate, glass beads (acid washed, 100) were dispensed using a pre-filled custom-made plate releasing approximately 100 mg beads/well, followed by centrifugation (0.5 min, 4°C, 1000 g). Then, 200 µl of freshly prepared 7M urea, 0.1M ammonium bicarbonate (ABC) were added to each well. Plates were sealed using Cap Mats and cells were lysed by bead milling with a Genogrinder (MiniG, SPEX) for 5 min at 1500 rpm, followed by quick centrifugation (1 min, 4°C, 3000 g). Samples were then processed as previously described^32^ on a Biomek i7 pipetting robot. To this end, 20 µl 5 mM DTT was added to each well, mixed, and shortly centrifuged and incubated for 1h at 30°C. Sample was left at room temperature for 15 min, and 20 µl 5 mM DTT was added, mixed, and briefly centrifuged, and incubated for 30 min in the dark at room temperature. Reduced and alkylated samples were then diluted with 1000 µl 0.1M ABC, mixed and centrifuged shortly, and 500 µl diluted lysate was transferred to a plate containing 2 µg trypsin/LysC per well, and incubated for 17h at 37°C. The digest was stopped by addition of 25 µl 20% formic acid, and purified using solid-phase extraction in 96-well format. Plates were conditioned with 200µl methanol (centrifuged at 50 g), washed twice 200 µl with 50% acetonitrile/water (centrifuged at 50 g), equilibrated thrice with 3% acetonitrile/0.1% formic acid in water (centrifuged at 50 g, 80 g, 100g). 500 µl per well was loaded (centrifuged at 100 g) and washed thrice with 200µl 3% acetonitrile/0.1% formic acid in water (centrifuged at 100 g), followed by another centrifugation step at 180 g. Peptides were eluted in two steps with 120 µl and 150 µl 50% acetonitrile/water, and dried to completeness in a vacuum concentrator. Samples were then redissolved in 3% acetonitrile/0.1% formic acid, and ready for analysis.

#### Liquid chromatography/mass spectrometry

For proteomics, digested peptides were separated on a high-flow chromatographic gradient and recorded by mass spectrometry using Scanning SWATH^23^ on an Agilent Infinity II HPLC combined with a SCIEX 6600 TripleTOF platform. Five micrograms of sample were injected onto a reverse phase HPLC column (Luna®Omega 1.6µm C18 100A, 30 × 2.1 mm, Phenomenex) and resolved by gradient elution at column temperature of 30⁰C with 0.1% formic acid in water (Solvent A) and 0.1% formic acid in acetonitrile (Solvent B). All solvents were of LC-MS grade. The gradient separation was at a flow rate of 0. 8 ml/min flow with the steps 0 min (1 % B), 0.1 min (5% B), 2.65 min (32% B), 3 min (40% B), followed by wash steps with 1.2ml/min flow at 3.5 min (80% B) to 3.7 min (80% B), and column equilibration with 1 ml/min flow from 3.8 min (1% B) to 4.8 min (1% B). For mass spectrometry analysis, the scanning SWATH precursor isolation window was 10 m/z, the bin size was set to 20% of the window size, the cycle time was 0.41 s, the precursor range was set to 400 - 900 m/z, the fragment range to 100 - 1500 m/z as previously described in Messner et al. ^23^. A Sciex IonDrive TurboV source was used with ion source gas 1 (nebulizer gas), ion source gas 2 (heater gas) and curtain gas set to 50 psi, 40 psi and 35 psi, respectively. The source temperature and ion spray voltage were set to 450⁰C and 5500 V, respectively.

For validation of reconstructed strains from batch 3, proteome samples were analyzed on a ZenoTOF 7600 system mass spectrometer (SCIEX), coupled to a 1290 Infinity II LC (Agilent). Prior to MS analysis, peptides were chromatographically separated on a Phenomenex Luna®Omega column (1.6μm C18 100A, 30 × 2.1 mm) heated to 50°C, using a flow rate of 0.5 ml / min where mobile phase A & B are 0.1% formic acid in water and 0.1% formic acid in acetonitrile, respectively. The gradient program was as follows: 1% to 36% B in 5 min, increase to 80% B at 0.8 mL over 0.5 min, which was maintained for 0.2 min and followed by equilibration with starting conditions for 2 min. For data independent acquisition Zeno SWATH MS/MS acquisition scheme was used with 80 variable size windows and 13 ms accumulation time. Ion source parameters were set to: Ion source gas 1 and 2 were set as 60 and 65 psi respectively; curtain gas 55, CAD gas 7 and source temperature at 600°C; Spray voltage was set at 4000V.

#### CRISPR genome editing

Genome editing was conducted as described in^35^. Briefly, yeast transformed with appropriate CRISPEY gene editing plasmids were induced for editing in galactose, quenched on YPD, and single colonies lacking the editing plasmid were isolated by selection on 5-FOA. Candidate edited strains were genotyped by PCR amplification of the relevant locus followed by Sanger sequencing.

### Quantification and statistical analysis

#### DIA-NN quantification and data processing

Mass spectrometry data was processed using an experimentally derived gas-phase fractionation spectral library using the DIA-NN software^26^ (version 1.8) with MS1 mass accuracy of 1.2×10^-5^, MS2 mass accuracy of 2×10^-5^, and a scan window radius of 6. Blanks and poorly growing samples (Z-scored OD_600_ < −2.5) were excluded, as were non-proteotypic precursors and entries with either Global.Q.Value, Global.PG.Q.Value, Q.Value, or PG.Q.Value > 0.01. Precursors were filtered to those occurring in > 80% of samples and those with CV > 0.3 in quality control injections were excluded. To account for plate effects, the plate-wise median for each precursor was adjusted to the grand median across all samples. Protein groups were quantified using maxLFQ^82^ in the DIA-NN R package^26^; a total of 1,225 proteins were identified across 1,042 samples. Proteomic differences were similarly distributed between high- and low-abundance proteins; with the exception of the lowest abundance fraction; their higher variance may be due in part to technical variability. After batch correction, we obtained proteomes with a median technical coefficient of variation (CV) on proteins of ∼ 11.0%. The proteomes contained few missing values, allowing stringent filtering: peptides shared across at least 80% of samples quantified 1,225 proteins, with an average of just 2.3% missing values [**Supplemental Table S2**].

#### Simulations and power calculations

We estimated the sensitivity of our pQTL mapping approach using *in silico* simulated protein abundance traits. Briefly, we generated simulated protein abundance vectors and performed pQTL mapping across a range of key parameters, including the number of F_6_ progeny used and the number of underlying pQTLs per protein. Summary results of these simulations can be found in **Fig. S7C**. Based on these data, we conducted our mapping experiment with the greatest possible number of F_6_ haploid isolates that were auxotrophic only for uracil, to maximize our sensitivity to pQTLs of modest effect.

#### Heritability estimates

We estimated broad-sense protein abundance heritability separately for each haploid parent control (RM11 and YJM975) using a linear mixed effect model that accounted for the harvest optical density (OD_600_) of each control sample. These estimates accorded well between the parental controls [**Fig. S7D**].

#### Genetic mapping

Genetic mapping was conducted essentially as in^22^ using protein abundance as the quantitative trait. Protein group abundance estimates from DIA-NN and maxLFQ were normalized to mean 0 and standard deviation 1, and we appended to the haploid genotype matrix a ‘pseudo-genotype’ representing the harvest OD_600_ of each sample (see also **Fig. S1H**). Following coarse mapping of pQTLs by stepwise selection, fine mapping of pQTNs was performed by ANOVA as described previously^22^. False discovery rate was estimated per-protein by 100 permutations of real abundance data; the empirical *p* value cutoffs were set to achieve ∼ 10% FDR. This procedure was conducted for the entire genotype matrix in the so-called ‘global’ mapping. In parallel, we conducted ‘local’ mapping that only considered loci within 10 markers of the ORF encoding the protein in question. Empirically, we found that putative *cis*-acting pQTL effects accorded well between the global and local approaches [**Fig. S1I**]; the analyses in the paper are based on the global analysis.

#### Mutation simulations and protein structure analysis

Simulations of all possible missense variants were conducted on the basis of the S288C reference genome R64. Briefly, we generated *in silico* all possible single-nucleotide changes to all S288C ORFs and categorized these as missense or synonymous and as transitions or transversions. Allele frequencies for extant variants were determined with reference to the 1,002 Yeast Genomes genotype matrix.

Predicted protein structures of all *S. cerevisiae* S288C ORFs were retrieved from https://alphafold.ebi.ac.uk/download#proteomes-section. Each ORF was analyzed with DSSP^83^ as well as using custom code to calculate the number of neighboring alpha-carbons. Based on these analyses, we annotated each possible missense SNP generated above with these structural parameters.

#### Phenotypic mapping

Phenotype data for ∼15,000 F_6_ diploid isolates from the RM11 x YM975 cross grown in various environmental conditions were released previously as part of our study of the effects of Hsp90 on the genotype-to-phenotype map^34^. Here, we reanalyzed the control dataset (without Hsp90 inhibition) from that study to identify QTLs and QTNs for growth under stress. Genetic mapping was conducted essentially as described above and previously^22^; complete mapping results can be found in **Supplemental Table S7**.

## References

1. Landrum, M.J., Lee, J.M., Riley, G.R., Jang, W., Rubinstein, W.S., Church, D.M., and Maglott, D.R. (2014). ClinVar: public archive of relationships among sequence variation and human phenotype. Nucleic Acids Res. 42, D980–D985.

2. Gudmundsson, S., Singer-Berk, M., Watts, N.A., Phu, W., Goodrich, J.K., Solomonson, M., Genome Aggregation Database Consortium, Rehm, H.L., MacArthur, D.G., and O’Donnell-Luria, A. (2022). Variant interpretation using population databases: Lessons from gnomAD. Hum. Mutat. 43, 1012–1030.

3. Leiding, J.W., Vogel, T.P., Santarlas, V.G.J., Mhaskar, R., Smith, M.R., Carisey, A., Vargas-Hernández, A., Silva-Carmona, M., Heeg, M., Rensing-Ehl, A., et al. (2023). Monogenic early-onset lymphoproliferation and autoimmunity: Natural history of STAT3 gain-of-function syndrome. J. Allergy Clin. Immunol. 151, 1081–1095.

4. GTEx Consortium (2017). Genetic effects on gene expression across human tissues. Nature 550, 204–213.

5. Wagner, N., Çelik, M.H., Hölzlwimmer, F.R., Mertes, C., Prokisch, H., Yépez, V.A., and Gagneur, J. (2023). Aberrant splicing prediction across human tissues. Nat. Genet. 55, 861– 870.

6. Foss, E.J., Radulovic, D., Shaffer, S.A., Ruderfer, D.M., Bedalov, A., Goodlett, D.R., and Kruglyak, L. (2007). Genetic basis of proteome variation in yeast. Nat. Genet. 39, 1369– 1375.

7. Damerval, C., Maurice, A., Josse, J.M., and de Vienne, D. (1994). Quantitative trait loci underlying gene product variation: a novel perspective for analyzing regulation of genome expression. Genetics 137, 289–301.

8. Suhre, K., McCarthy, M.I., and Schwenk, J.M. (2021). Genetics meets proteomics: perspectives for large population-based studies. Nat. Rev. Genet. 22, 19–37.

9. Pietzner, M., Wheeler, E., Carrasco-Zanini, J., Cortes, A., Koprulu, M., Wörheide, M.A., Oerton, E., Cook, J., Stewart, I.D., Kerrison, N.D., et al. (2021). Mapping the proteo-genomic convergence of human diseases. Science 374, eabj1541.

10. Brem, R.B., Yvert, G., Clinton, R., and Kruglyak, L. (2002). Genetic Dissection of Transcriptional Regulation in Budding Yeast. Science 296, 752–755.

11. Yvert, G., Brem, R.B., Whittle, J., Akey, J.M., Foss, E., Smith, E.N., Mackelprang, R., and Kruglyak, L. (2003). *Trans*-acting regulatory variation in *Saccharomyces cerevisiae* and the role of transcription factors. Nat. Genet. 35, 57–64.

12. Albert, F.W., Bloom, J.S., Siegel, J., Day, L., and Kruglyak, L. (2018). Genetics of trans-regulatory variation in gene expression. eLife Sciences 7, e35471.

13. Picotti, P., Clément-Ziza, M., Lam, H., Campbell, D.S., Schmidt, A., Deutsch, E.W., Röst, H., Sun, Z., Rinner, O., Reiter, L., et al. (2013). A complete mass-spectrometric map of the yeast proteome applied to quantitative trait analysis. Nature 494, 266–270.

14. Albert, F.W., Treusch, S., Shockley, A.H., Bloom, J.S., and Kruglyak, L. (2014). Genetics of single-cell protein abundance variation in large yeast populations. Nature 506, 494–497.

15. Parts, L., Liu, Y.-C., Tekkedil, M.M., Steinmetz, L.M., Caudy, A.A., Fraser, A.G., Boone, C., Andrews, B.J., and Rosebrock, A.P. (2014). Heritability and genetic basis of protein level variation in an outbred population. Genome Res. 24, 1363–1370.

16. Vande Zande, P., Hill, M.S., and Wittkopp, P.J. (2022). Pleiotropic effects of trans-regulatory mutations on fitness and gene expression. Science 377, 105–109.

17. Schadt, E.E., Monks, S.A., Drake, T.A., Lusis, A.J., Che, N., Colinayo, V., Ruff, T.G., Milligan, S.B., Lamb, J.R., Cavet, G., et al. (2003). Genetics of gene expression surveyed in maize, mouse and man. Nature 422, 297–302.

18. Grossbach, J., Gillet, L., Clément-Ziza, M., Schmalohr, C.L., Schubert, O.T., Schütter, M., Mawer, J.S.P., Barnes, C.A., Bludau, I., Weith, M., et al. (2022). The impact of genomic variation on protein phosphorylation states and regulatory networks. Mol. Syst. Biol. 18, e10712.

19. Schubert, O.T., Bloom, J.S., Sadhu, M.J., and Kruglyak, L. (2022). Genome-wide base editor screen identifies regulators of protein abundance in yeast. 10.1101/2022.03.09.483657.

20. Peter, J., Chiara, M.D., Friedrich, A., Yue, J.-X., Pflieger, D., Bergström, A., Sigwalt, A., Barre, B., Freel, K., Llored, A., et al. (2018). Genome evolution across 1,011 Saccharomyces cerevisiae isolates. Nature, 1.

21. Caudal, É., Loegler, V., Dutreux, F., Vakirlis, N., Teyssonnière, É., Caradec, C., Friedrich, A., Hou, J., and Schacherer, J. (2024). Pan-transcriptome reveals a large accessory genome contribution to gene expression variation in yeast. Nat. Genet. 56, 1278–1287.

22. She, R., and Jarosz, D.F. (2018). Mapping Causal Variants with Single-Nucleotide Resolution Reveals Biochemical Drivers of Phenotypic Change. Cell 172, 478–490.e15.

23. Messner, C.B., Demichev, V., Bloomfield, N., Yu, J.S.L., White, M., Kreidl, M., Egger, A.- S., Freiwald, A., Ivosev, G., Wasim, F., et al. (2021). Ultra-fast proteomics with Scanning SWATH. Nat. Biotechnol. 39, 846–854.

24. McCullough, M.J., Clemons, K.V., Farina, C., McCusker, J.H., and Stevens, D.A. (1998). Epidemiological investigation of vaginal Saccharomyces cerevisiae isolates by a genotypic method. J. Clin. Microbiol. 36, 557–562.

25. Török, T., Mortimer, R.K., Romano, P., Suzzi, G., and Polsinelli, M. (1996). Quest for wine yeasts—An old story revisited. J. Ind. Microbiol. Biotechnol. 17, 303–313.

26. Demichev, V., Messner, C.B., Vernardis, S.I., Lilley, K.S., and Ralser, M. (2019). DIA-NN: neural networks and interference correction enable deep proteome coverage in high throughput. Nat. Methods 17, 41–44.

27. Liti, G., Carter, D.M., Moses, A.M., Warringer, J., Parts, L., James, S.A., Davey, R.P., Roberts, I.N., Burt, A., Koufopanou, V., et al. (2009). Population genomics of domestic and wild yeasts. Nature 458, 337–341.

28. Huang, Q., Szklarczyk, D., Wang, M., Simonovic, M., and von Mering, C. (2023). PaxDb 5.0: Curated Protein Quantification Data Suggests Adaptive Proteome Changes in Yeasts. Mol. Cell. Proteomics 22, 100640.

29. Rolland, T., and Dujon, B. (2011). Yeasty clocks: dating genomic changes in yeasts. C. R. Biol. 334, 620–628.

30. Messner, C.B., Demichev, V., Muenzner, J., Aulakh, S.K., Barthel, N., Röhl, A., Herrera-Domínguez, L., Egger, A.-S., Kamrad, S., Hou, J., et al. (2023). The proteomic landscape of genome-wide genetic perturbations. Cell 186, 2018–2034.e21.

31. Hahne, K., Haucke, V., Ramage, L., and Schatz, G. (1994). Incomplete arrest in the outer membrane sorts NADH-cytochrome b5 reductase to two different submitochondrial compartments. Cell 79, 829–839.

32. Muenzner, J., Trébulle, P., Agostini, F., Zauber, H., Messner, C.B., Steger, M., Kilian, C., Lau, K., Barthel, N., Lehmann, A., et al. (2024). Natural proteome diversity links aneuploidy tolerance to protein turnover. Nature 630, 149–157.

33. Teyssonnière, E.M., Trébulle, P., Muenzner, J., Loegler, V., Ludwig, D., Amari, F., Mülleder, M., Friedrich, A., Hou, J., Ralser, M., et al. (2024). Species-wide quantitative transcriptomes and proteomes reveal distinct genetic control of gene expression variation in yeast. Proc. Natl. Acad. Sci. U. S. A. 121, e2319211121.

34. Jakobson, C.M., Aguilar-Rodríguez, J., and Jarosz, D.F. (2023). Hsp90 shapes adaptation by controlling the fitness consequences of regulatory variation. bioRxiv. 10.1101/2023.10.30.564848.

35. Sharon, E., Chen, S.-A.A., Khosla, N.M., Smith, J.D., Pritchard, J.K., and Fraser, H.B. (2018). Functional Genetic Variants Revealed by Massively Parallel Precise Genome Editing. Cell. 10.1016/j.cell.2018.08.057.

36. Zambelli, F., Pesole, G., and Pavesi, G. (2009). Pscan: finding over-represented transcription factor binding site motifs in sequences from co-regulated or co-expressed genes. Nucleic Acids Res. 37, W247–W252.

37. Tanaka, K., Matsumoto, K., and Toh-E, A. (1989). IRA1, an Inhibitory Regulator of the RAS-Cyclic AMP Pathway in Saccharomyces cerevisiae. Mol. Cell. Biol. 9, 757–768.

38. Tanaka, K., Lin, B.K., Wood, D.R., and Tamanoi, F. (1991). IRA2, an upstream negative regulator of RAS in yeast, is a RAS GTPase-activating protein. Proceedings of the National Academy of Sciences 88, 468–472.

39. Sass, P., Field, J., Nikawa, J., Toda, T., and Wigler, M. (1986). Cloning and characterization of the high-affinity cAMP phosphodiesterase of Saccharomyces cerevisiae. Proceedings of the National Academy of Sciences 83, 9303–9307.

40. D’Souza, C.A., and Heitman, J. (2001). Conserved cAMP signaling cascades regulate fungal development and virulence. FEMS Microbiol. Rev. 25, 349–364.

41. Molinet, J., Navarrete, J.P., Villarroel, C.A., Villarreal, P., Sandoval, F.I., Nespolo, R.F., Stelkens, R., and Cubillos, F.A. (2024). Wild Patagonian yeast improve the evolutionary potential of novel interspecific hybrid strains for lager brewing. PLoS Genet. 20, e1011154.

42. Hogan, D.A., and Sundstrom, P. (2009). The Ras/cAMP/PKA signaling pathway and virulence in Candida albicans. Future Microbiol. 4, 1263–1270.

43. Pedruzzi, I., Bürckert, N., Egger, P., and De Virgilio, C. (2000). Saccharomyces cerevisiae Ras/cAMP pathway controls post-diauxic shift element-dependent transcription through the zinc finger protein Gis1. EMBO J. 19, 2569–2579.

44. Kemmeren, P., Sameith, K., van de Pasch, L.A.L., Benschop, J.J., Lenstra, T.L., Margaritis, T., O’Duibhir, E., Apweiler, E., van Wageningen, S., Ko, C.W., et al. (2014). Large-scale genetic perturbations reveal regulatory networks and an abundance of gene-specific repressors. Cell 157, 740–752.

45. Orr, H.A. (1998). Testing Natural Selection vs. Genetic Drift in Phenotypic Evolution Using Quantitative Trait Locus Data. Genetics 149, 2099–2104.

46. Henikoff, S., and Henikoff, J.G. (1992). Amino acid substitution matrices from protein blocks. Proc. Natl. Acad. Sci. U. S. A. 89, 10915–10919.

47. Schymkowitz, J., Borg, J., Stricher, F., Nys, R., Rousseau, F., and Serrano, L. (2005). The FoldX web server: an online force field. Nucleic Acids Res. 33, W382–W388.

48. Jumper, J., Evans, R., Pritzel, A., Green, T., Figurnov, M., Ronneberger, O., Tunyasuvunakool, K., Bates, R., Žídek, A., Potapenko, A., et al. (2021). Highly accurate protein structure prediction with AlphaFold. Nature 596, 583–589.

49. Costanzo, M., Baryshnikova, A., Bellay, J., Kim, Y., Spear, E.D., Sevier, C.S., Ding, H., Koh, J.L.Y., Toufighi, K., Mostafavi, S., et al. (2010). The Genetic Landscape of a Cell. Science 327, 425–431.

50. Giaever, G., Chu, A.M., Ni, L., Connelly, C., Riles, L., Véronneau, S., Dow, S., Lucau-Danila, A., Anderson, K., André, B., et al. (2002). Functional profiling of the *Saccharomyces cerevisiae* genome. Nature 418, 387–391.

51. Ghaemmaghami, S., Huh, W.-K., Bower, K., Howson, R.W., Belle, A., Dephoure, N., O’Shea, E.K., and Weissman, J.S. (2003). Global analysis of protein expression in yeast. Nature 425, 737–741.

52. Rodríguez, A., De La Cera, T., Herrero, P., and Moreno, F. (2001). The hexokinase 2 protein regulates the expression of the GLK1, HXK1 and HXK2 genes of Saccharomyces cerevisiae. Biochem. J 355, 625–631.

53. Umekawa, M., Hamada, K., Isono, N., and Karita, S. (2020). The Emi2 Protein of Saccharomyces cerevisiae is a Hexokinase Expressed under Glucose Limitation. J. Appl. Glycosci. 67, 103–109.

54. Muenzner, J., Trébulle, P., Agostini, F., Messner, C.B., Steger, M., Lehmann, A., Caudal, E., Egger, A.-S., Amari, F., Barthel, N., et al. (2022). The natural diversity of the yeast proteome reveals chromosome-wide dosage compensation in aneuploids. Preprint, 10.1101/2022.04.06.487392 10.1101/2022.04.06.487392.

55. Meldal, B.H.M., Perfetto, L., Combe, C., Lubiana, T., Ferreira Cavalcante, J.V., Bye-A-Jee, H., Waagmeester, A., del-Toro, N., Shrivastava, A., Barrera, E., et al. (2022). Complex Portal 2022: new curation frontiers. Nucleic Acids Res. 50, D578–D586.

56. Szklarczyk, D., Kirsch, R., Koutrouli, M., Nastou, K., Mehryary, F., Hachilif, R., Gable, A.L., Fang, T., Doncheva, N.T., Pyysalo, S., et al. (2023). The STRING database in 2023: protein-protein association networks and functional enrichment analyses for any sequenced genome of interest. Nucleic Acids Res. 51, D638–D646.

57. Costanzo, M., VanderSluis, B., Koch, E.N., Baryshnikova, A., Pons, C., Tan, G., Wang, W., Usaj, M., Hanchard, J., Lee, S.D., et al. (2016). A global genetic interaction network maps a wiring diagram of cellular function. Science 353, aaf1420.

58. Meldal, B.H.M., Bye-A-Jee, H., Gajdoš, L., Hammerová, Z., Horácková, A., Melicher, F., Perfetto, L., Pokorný, D., Lopez, M.R., Türková, A., et al. (2019). Complex Portal 2018: extended content and enhanced visualization tools for macromolecular complexes. Nucleic Acids Res. 47, D550–D558.

59. Oughtred, R., Rust, J., Chang, C., Breitkreutz, B.-J., Stark, C., Willems, A., Boucher, L., Leung, G., Kolas, N., Zhang, F., et al. (2021). The BioGRID database: A comprehensive biomedical resource of curated protein, genetic, and chemical interactions. Protein Sci. 30, 187–200.

60. Esnault, Y., Feldheim, D., Blondel, M.O., Schekman, R., and Képès, F. (1994). SSS1 encodes a stabilizing component of the Sec61 subcomplex of the yeast protein translocation apparatus. J. Biol. Chem. 269, 27478–27485.

61. Galello, F., Moreno, S., and Rossi, S. (2014). Interacting proteins of protein kinase A regulatory subunit in Saccharomyces cerevisiae. J. Proteomics 109, 261–275.

62. Dancis, A., Klausner, R.D., Hinnebusch, A.G., and Barriocanal, J.G. (1990). Genetic evidence that ferric reductase is required for iron uptake in Saccharomyces cerevisiae. Mol. Cell. Biol. 10, 2294–2301.

63. Vowinckel, J., Hartl, J., Marx, H., Kerick, M., Runggatscher, K., Keller, M.A., Mülleder, M., Day, J., Weber, M., Rinnerthaler, M., et al. (2021). The metabolic growth limitations of petite cells lacking the mitochondrial genome. Nat Metab 3, 1521–1535.

64. Mattoon, J.R., Caravajal, E., and Guthrie, D. (1990). Effects of hap mutations on heme and cytochrome formation in yeast. Curr. Genet. 17, 179–183.

65. McNabb, D.S., and Pinto, I. (2005). Assembly of the Hap2p/Hap3p/Hap4p/Hap5p-DNA complex in Saccharomyces cerevisiae. Eukaryot. Cell 4, 1829–1839.

66. Fragiadakis, G.S., Tzamarias, D., and Alexandraki, D. (2004). Nhp6 facilitates Aft1 binding and Ssn6 recruitment, both essential for FRE2 transcriptional activation. EMBO J. 23, 333–342.

67. Ramos-Alonso, L., Romero, A.M., Martínez-Pastor, M.T., and Puig, S. (2020). Iron Regulatory Mechanisms in Saccharomyces cerevisiae. Front. Microbiol. 11, 582830.

68. Karst, F., and Lacroute, F. (1977). Ergosterol biosynthesis inSaccharomyces cerevisiae. Mol. Gen. Genet. 154, 269–277.

69. Jakobson, C.M., She, R., and Jarosz, D.F. (2019). Pervasive function and evidence for selection across standing genetic variation in S. cerevisiae. Nat. Commun. 10, 1222.

70. Jellen-Ritter, A.S., and Kern, W.V. (2001). Enhanced expression of the multidrug efflux pumps AcrAB and AcrEF associated with insertion element transposition in Escherichia coli mutants Selected with a fluoroquinolone. Antimicrob. Agents Chemother. 45, 1467– 1472.

71. Bloom, J.S., Ehrenreich, I.M., Loo, W.T., Lite, T.-L.V., and Kruglyak, L. (2013). Finding the sources of missing heritability in a yeast cross. Nature 494, 234–237.

72. Boyle, E.A., Li, Y.I., and Pritchard, J.K. (2017). An Expanded View of Complex Traits: From Polygenic to Omnigenic. Cell 169, 1177–1186.

73. Rieseberg, L.H., Archer, M.A., and Wayne, R.K. (1999). Transgressive segregation, adaptation and speciation. Heredity 83 ( *Pt 4*), 363–372.

74. Albert, F.W., and Kruglyak, L. (2015). The role of regulatory variation in complex traits and disease. Nat. Rev. Genet. 16, 197–212.

75. Kern, A.D., and Hahn, M.W. (2018). The Neutral Theory in Light of Natural Selection. Mol. Biol. Evol. 35, 1366–1371.

76. Lemke, O., Heineike, B.M., Viknander, S., Cohen, N., Steenwyk, J.L., Spranger, L., Li, F., Agostini, F., Lee, C.T., Aulakh, S.K., et al. (2024). The Role of Metabolism in Shaping Enzyme Structures Over 400 Million Years of Evolution. bioRxiv, 2024.05.27.596037. 10.1101/2024.05.27.596037.

77. Besse, S., Sakaguchi, T., Gauthier, L., Sahaf, Z., Péloquin, O., Gonzalez, L., Castellanos-Girouard, X., Koçatug, N., Matta, C., Hussin, J.G., et al. (2023). Genetic landscape of an in vivo protein interactome. bioRxiv, 2023.12.14.571726. 10.1101/2023.12.14.571726.

78. Fischer, J.D., Holliday, G.L., Rahman, S.A., and Thornton, J.M. (2010). The structures and physicochemical properties of organic cofactors in biocatalysis. J. Mol. Biol. 403, 803–824.

79. Piazza, I., Kochanowski, K., Cappelletti, V., Fuhrer, T., Noor, E., Sauer, U., and Picotti, P. (2018). A map of protein-metabolite interactions reveals principles of chemical communication. Cell 172, 358–372.e23.

80. Lutz, S., Van Dyke, K., Feraru, M.A., and Albert, F.W. (2022). Multiple epistatic DNA variants in a single gene affect gene expression in trans. Genetics 220. 10.1093/genetics/iyab208.

81. Manolio, T.A., Collins, F.S., Cox, N.J., Goldstein, D.B., Hindorff, L.A., Hunter, D.J., McCarthy, M.I., Ramos, E.M., Cardon, L.R., Chakravarti, A., et al. (2009). Finding the missing heritability of complex diseases. Nature 461, 747–753.

82. Cox, J., Hein, M.Y., Luber, C.A., Paron, I., Nagaraj, N., and Mann, M. (2014). Accurate Proteome-wide Label-free Quantification by Delayed Normalization and Maximal Peptide Ratio Extraction, Termed MaxLFQ*. Mol. Cell. Proteomics 13, 2513–2526.

83. Joosten, R.P., te Beek, T.A.H., Krieger, E., Hekkelman, M.L., Hooft, R.W.W., Schneider, R., Sander, C., and Vriend, G. (2010). A series of PDB related databases for everyday needs. Nucleic Acids Res. 39, D411–D419.

84. Kabsch, W., and Sander, C. (1983). Dictionary of protein secondary structure: pattern recognition of hydrogen-bonded and geometrical features. Biopolymers 22, 2577–2637.

85. Lawless, C., Holman, S.W., Brownridge, P., Lanthaler, K., Harman, V.M., Watkins, R., Hammond, D.E., Miller, R.L., Sims, P.F.G., Grant, C.M., et al. (2016). Direct and Absolute Quantification of over 1800 Yeast Proteins via Selected Reaction Monitoring. Mol. Cell. Proteomics 15, 1309–1322.

86. Srivastava, A.P., Luo, M., Zhou, W., Symersky, J., Bai, D., Chambers, M.G., Faraldo-Gómez, J.D., Liao, M., and Mueller, D.M. (2018). High-resolution cryo-EM analysis of the yeast ATP synthase in a lipid membrane. Science 360. 10.1126/science.aas9699.

