## Supplemental Information for "A genome-to-proteome atlas charts natural variants controlling proteome diversity and forecasts their fitness effects"

To accompany Jakobson*, Hartl*, *et al.*

**Contents:**

Key Resources Table

Figures S1-S7

Table S6

**Key Resources Table**

| Reagent or resource | Source | Identifier |
|---|---|---|
| *Chemicals* | | |
| Yeast nitrogen base | RPI | Y20040 |
| Glucose | Fisher | D16-3 |
| Uracil | Sigma | U0750 |
| Agar | IBI | IB49170 |
| Fluconazole | TCI | F0677 |
| Ethanol (95%) | Fisher | 04-355-226 |
| Water (LC-MS grade) | Fisher Scientific | Cat 10509404; CAS: 7732-18-5 |
| Acetonitrile (LC-MS grade) | Fisher Scientific | Cat 10489553; CAS: 75-05-8 |
| Methanol (LC-MS grade) | Fisher Scientific | Cat 10767665; CAS: 67-56-1 |
| Formic acid (LC-MS grade) | Fisher Scientific | Cat 5.33002; CAS: 64-18-6 |
| Dithiothreitol (≥99.5%) | Sigma Aldrich | Cat 43815; CAS: 3483-12-3 |
| Iodoacetamide (≥99%) | Sigma Aldrich | Cat I1149; CAS: 144-48-9 |
| Ammonium Bicarbonate | Sigma Aldrich | Cat 40867; CAS: 1066-33-7 |
| Urea | Sigma Aldrich | Cat 33247 |
| Trypsin/Lys-C (Mass Spec Grade) | Promega | V5072 |
| Solid-glass beads (borosilicate, diam. 4 mm) | Sigma Aldrich | Cat Z143936 |
| *Deposited data* | | |
| Mass spectrometry data | This study | Will be made available on the PRIDE archive upon acceptance |
| *Experimental models: Strains* | | |
| RM11 haploid | [34] | YDJ6649 |

| | | |
|---|---|---|
| YJM975 haploid | [34] | YDJ6635 |
| YJM975 *ERG11*$^{122014T>C}$ | [34] | YDJ8281 |
| YJM975 *ERG11*$^{Asn433Lys}$ | [34] | YDJ8436 |
| YJM975 *ERG11*$^{122014T>C; Asn433Lys}$ | [34] | YDJ8437 |
| RM11 *MCR1*$^{G>A}$ | This study | YDJ8524 |
| RM11 *NCP1*$^{A>T}$ | This study | YDJ8525 |
| RM11 *SER2*$^{G>A}$ | This study | YDJ8526 |
| RM11 *AAT2*$^{G>A}$ | This study | YDJ8527 |
| YJM975 *GCS1*$^{C>T}$ | This study | YDJ8528 |
| RM11 *IRA2*$^{G>A}$ | This study | YDJ8578 |
| YJM975 *IRA2*$^{A>G}$ | This study | YDJ8529 |
| *Oligonucleotides* | | |
| *MCR1* CRISPEY editing oligo<br>GAGTTACTGTCTGTTTTC<br>CTGTTACTTACTTTGTTG<br>ACGACAAGCAAGATGAC<br>CAAGACTTTGATGGTGA<br>AATTAGTTTCATCTCCAA<br>AGATTTTATTCAGGAGC<br>ATGTTCCAGGTCCAAAG<br>GAAACCCGTTTCTTCTGA<br>CGTAAGGGTGCGCACAA<br>GACTTTGATGGTGAAAT<br>GTTTCAGAGCTATGCTGG<br>AA | This study | CMJP697 |
| *NCP1* CRISPEY editing oligo<br>GAGTTACTGTCTGTTTTC<br>CTGGTCAACCCGCTATTG<br>TTCTCCAGCCAGCTTTTA<br>TCGTTTTGCATTTTTTTTT<br>CGGGCTGCTTTTCGTTCT<br>TCGAGGACAAACGCACC<br>TGTAAAGCTCAGAGGAA | This study | CMJP703 |

| | | |
|---|---|---|
| ACCCGTTTCTTCTGACGTAAGGGTGCGCAATCGTTTTGCATATTTTTTTGTTTCAGAGCTATGCTGGAA | | |
| *SER2* CRISPEY editing oligo<br>GAGTTACTGTCTGTTTTCCTTTCTTGGCTACACCGATGATGAAATATACAATAGACAATGAAGAAAATAATGATAGATAGATGTAATAGAGTTTCTTTTTAAAATTGTTTATTTAAACTGAGGAAACCCGTTTCTTCTGACGTAAGGGTGCGCAGACAATGAAGAAAATAATGAGTTTCAGAGCTATGCTGGAA | This study | CMJP721 |
| *AAT2* CRISPEY editing oligo<br>GAGTTACTGTCTGTTTTCCTAACTGCGTGGGTTTCTTCAAGTCGTTTAACCATTTGAGGAGTCAATCCTGTAAAGGAGAACATCCCGCATTGATTTACTATATGATCCCAGTTGCCAGGAAGGAAACCCGTTTCTTCTGACGTAAGGGTGCGCATTGAGGAGTCAACCCTGTAAGTTTCAGAGCTATGCTGGAA | This study | CMJP707 |
| *GCS1* CRISPEY editing oligo<br>GAGTTACTGTCTGTTTTCCTAATCCATACATTTCTTATTTGCACCAATCTTTTGCAATTGCAAAAGACGCCTACGGGTATCTGGGTCCACTTTCCAATCTGACATGCTCTATAATCCGCGAGGAAACCCGTTTCTTCTGACGTAAGGGTGCGCAGCAATTGCAAAAGACGCCTGGTTTCAGAGCTATGCTGGAA | This study | CMJP706 |

| | | |
|---|---|---|
| *IRA2* CRISPEY editing oligo (RM>YJM) GAGTTACTGTCTGTTTTCCTAAGTTCAATACAAGAACTTTGCAAATTTTACAAAATATGATCAGTCATGTTCATGGAAACATTCTAACGACTTTGAGTTCCTCGATTCTTCCCCGCCACAAGGAAACCCGTTTCTTCTGACGTAAGGGTGCGCAAGTATGATCAGTCATGTTCAGTTTCAGAGCTATGCTGGAA | This study | CMJP709 |
| *IRA2* CRISPEY editing oligo (YJM>RM) GAGTTACTGTCTGTTTTCCTAAGTTCAATACAAGAACTTTGCAAATTTTACAAAGTATGATCAGTCATGTTCATGGAAACATTCTAACGACTTTGAGTTCCTCGATTCTTCCCCGCCACAAGGAAACCCGTTTCTTCTGACGTAAGGGTGCGCAAATATGATCAGTCATGTTCAGTTTCAGAGCTATGCTGGAA | This study | CMJP710 |
| *Recombinant DNA* | | |
| CRISPEY editing plasmid: PDJ2318 | [34] | PDJ2318 |
| *Software and algorithms* | | |
| DSSP | [84] | https://swift.cmbi.umcn.nl/gv/dssp/index.html |
| DIA-NN | [26] | https://github.com/vdemichev/DiaNN |
| maxLFQ | [82] | https://rdrr.io/cran/iq/man/maxLFQ.html |
| *Other* | | |

| | | |
|---|---|---|
| Custom genetic mapping code | This study | https://github.com/cjakobson/pqtl-mapping |
| Genetic mapping dependencies | This study | https://www.dropbox.com/scl/fo/3xbcbe9ivwz8aahrlk137/APGxHor01S7jnNX3a1Yk3Og?rlkey=yx81ckrtaq8eb5pu80ggprjhs&dl=0 |
| Custom protein structure analysis code | This study | https://github.com/cjakobson/pop-gen-structure |
| Protein structure analysis dependencies | This study | https://www.dropbox.com/scl/fo/le2voq9djr79p2ehxqvs3/AGecke84_fLbzrsis2Ncpjk?rlkey=g5etvtwhay27j4y0sh6dh8dh7&dl=0 |

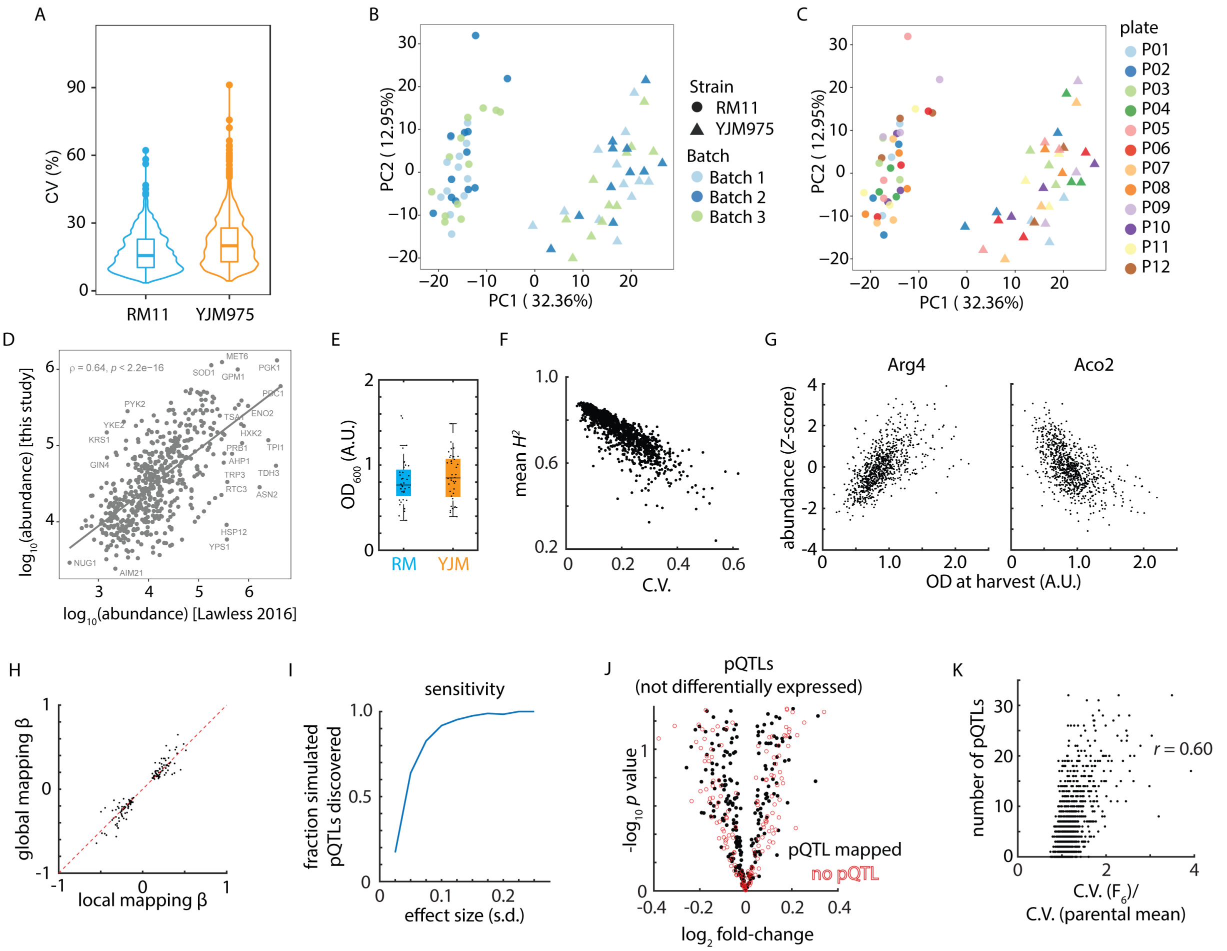


**Figure S1. To accompany Figure 1.** (A) Distribution of coefficients of variation (CVs) of all analyzed proteins (n=1225) from biological replicates of RM11 (n=36) and YJM975 (n=34) distributed and processed in 12 plates and measured in 3 LC-MS batches. (B-C) Variation according to principal components 1 and 2 of data described in (A) with only proteins without missing values (n=850) colored according to MS batch (B) or processing plate (C). (D) Median protein abundances from n=851 $F_6$ strains (ordinate) plotted and spearman correlated against absolute quantitative protein data from Lawless[85] (abscissa) for matching proteins (n=538). (E) Harvest $OD_{600}$ of RM and YJM biological replicate samples. (F) Mean broad-sense heritability (ordinate) as a function of technical variability (C.V.; abscissa). (G) Abundance of Arg4 (left) and Aco2 (right) as a function of harvest $OD_{600}$ amongst the $F_6$ progeny. (H) Genetic mapping coefficient (beta; units of st. dev.) from global (ordinate) and local (abscissa) approaches. Line of parity is shown in a red dashed line. (I) Genetic mapping sensitivity (fraction of simulated pQTLs discovered; ordinate) as a function of effect size (st. dev.; abscissa). Shown is the mean of $N = 100$ simulations of protein traits with 50 pQTLs. (J) Volcano plot illustrating $\log_2$ fold-change in protein abundance (abscissa) and Benjamini-Hochberg-corrected $t$ test $p$ value (ordinate) between the vineyard (RM) and clinical (YJM) parents highlighting proteins not differentially expressed between the parents. Closed symbols have a mapped pQTL; open symbols have no identified pQTLs. $n = 36 - 39$. (K) Number of pQTLs discovered (ordinate) as a function of normalized C.V. amongst the F6 progeny as compared to the mean C.V. in the parental isolates (abscissa).

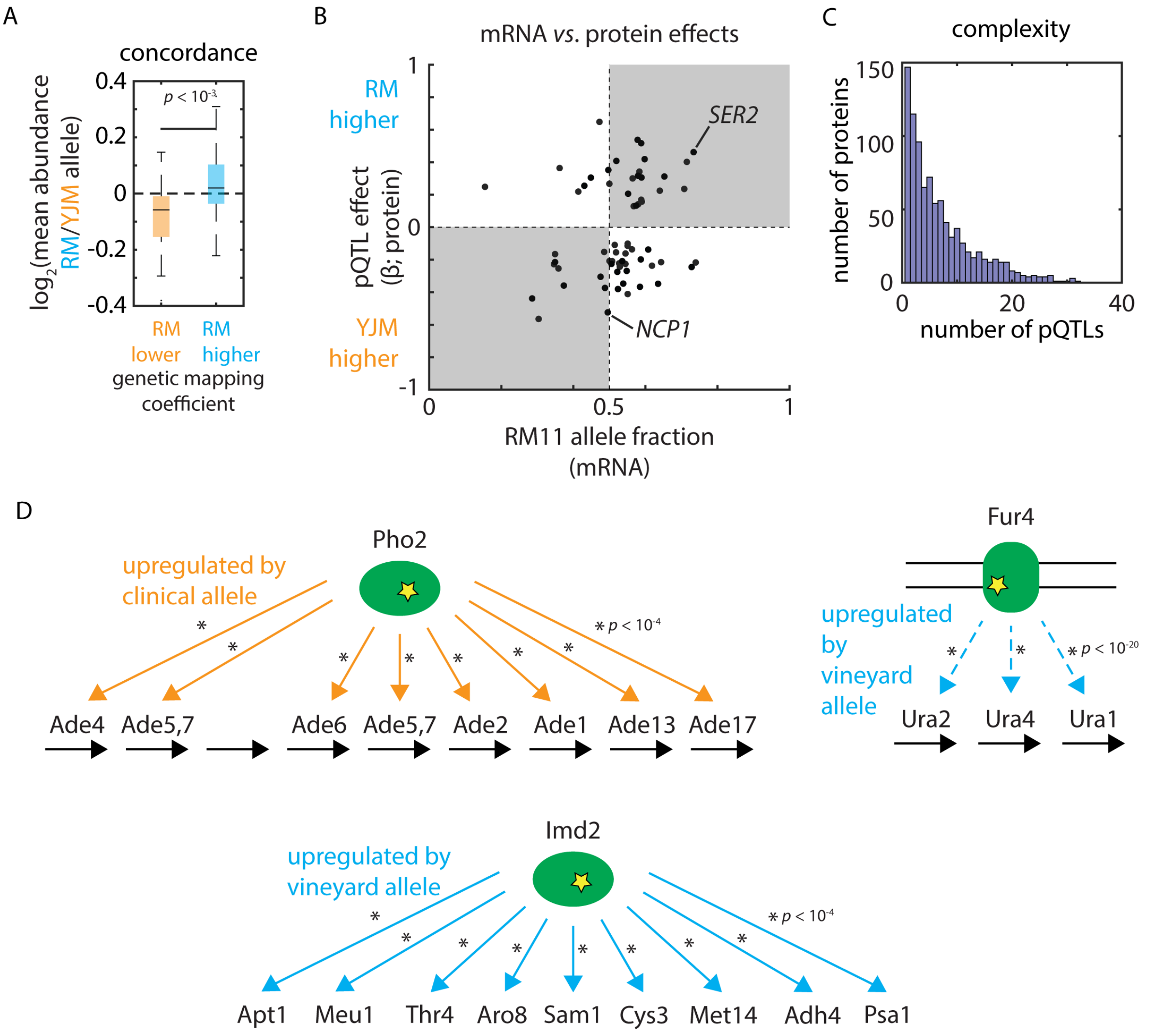


**Figure S2. To accompany Figure 2.** (A) Summary of replication across all *cis*-acting pQTLs we discovered; shown is $\log_2$ fold-change in mean protein abundance between 1,002 Yeast Genomes strains bearing the RM and YJM alleles in *cis*, divided by whether genetic mapping predicted the YJM or RM allele to exhibit higher protein level. $p$ value by two-sided $t$ test. (B) Predicted effect on protein level from genetic mapping (ordinate) and measured effect on mRNA level from allele-specific expression analysis (abscissa) for regulatory *cis*-pQTLs. Highlighted are *cis*-pQTNs selected for reconstruction. (C) Frequency of proteins (ordinate) as a function of the number of controlling pQTLs (abscissa). (D) Schematic of adenine biosynthetic pathway enzymes controlled by Pho2, uracil biosynthetic pathway enzymes controlled by Fur4, and metabolic enzymes controlled by Imd2.

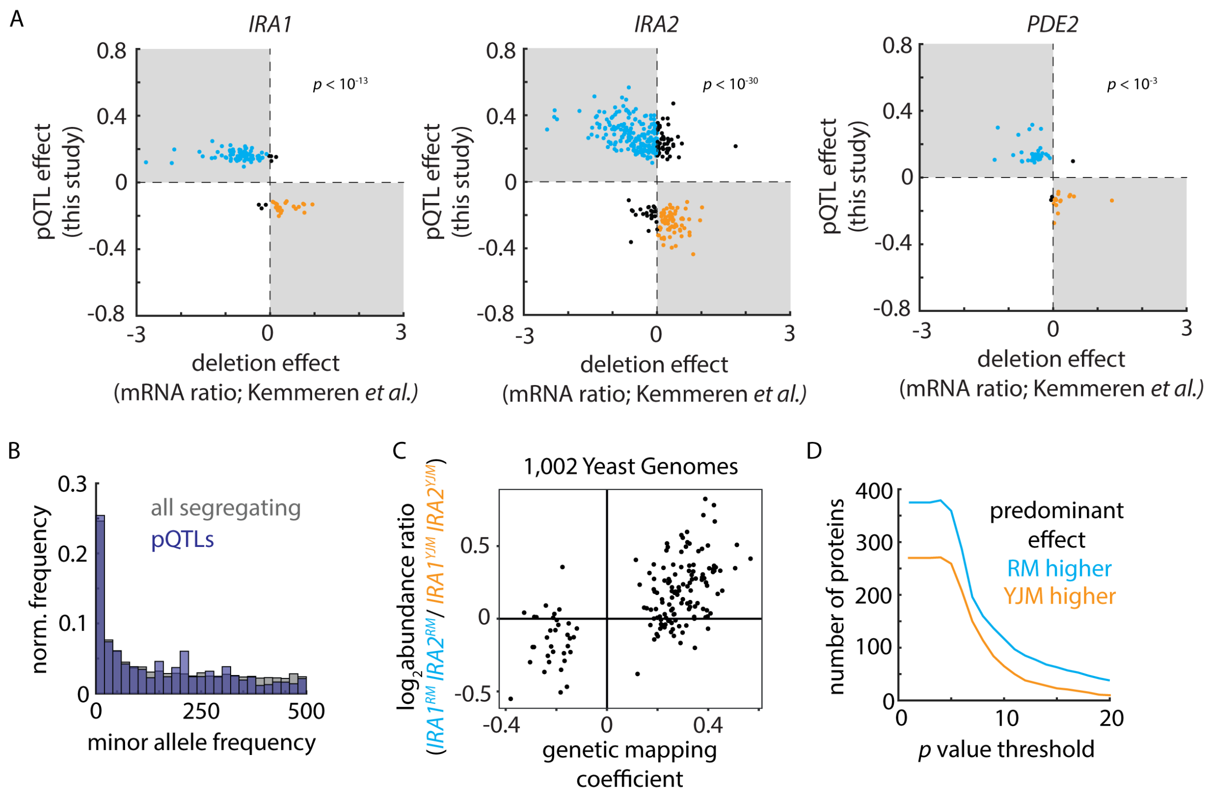


**Figure S3. To accompany Figure 3.** (A) Predicted effects of *trans* pQTNs from genetic mapping (ordinate) as a function of measured mRNA effects of deleting the corresponding gene (abscissa) [44] for *IRA1*, *IRA2*, and *PDE2*, as indicated. *p* values by *t* statistic. (B) Normalized frequency (ordinate) of minor allele frequencies amongst the 1,002 Genomes collection (absicssa) for pQTL variants (blue) and all segregating variation amongst the $F_6$ progeny (grey). (C) Relative abundance of Ira1 and Ira2 targets (ordinate; identified by pQTL mapping) in wild yeast proteomes bearing $IRA1^{RM}$ and $IRA2^{RM}$ alleles (n=5) as compared to strains with $IRA1^{YJM}$ and $IRA2^{YJM}$ (n=371) as a function of predicted effect from pQTL mapping (abscissa). (D) Number of proteins (ordinate) predominantly controlled by RM-higher *trans* pQTL alleles (blue) or YJM-higher *trans* pQTL alleles (orange) as a function of genetic mapping *p* value (abscissa).

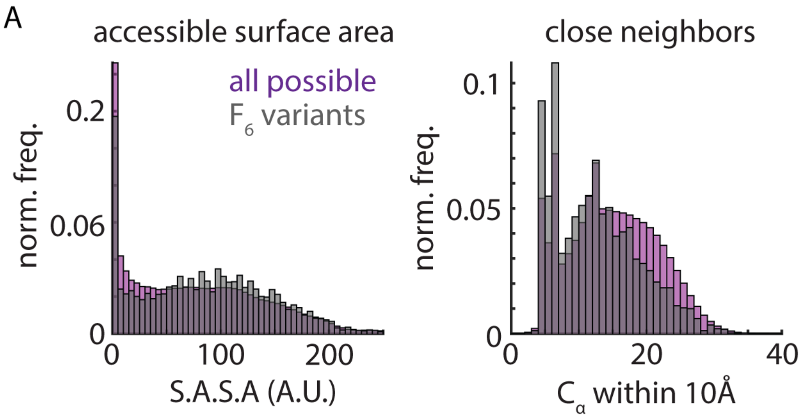


**Figure S4. To accompany Figure 4.** (A) Normalized frequencies of solvent-accessible surface area (left) and number of $C_\alpha$ within 10Å (right) for all possible missense SNPs (purple) and all missense variants segregating in the $F_6$ mapping panel (grey).

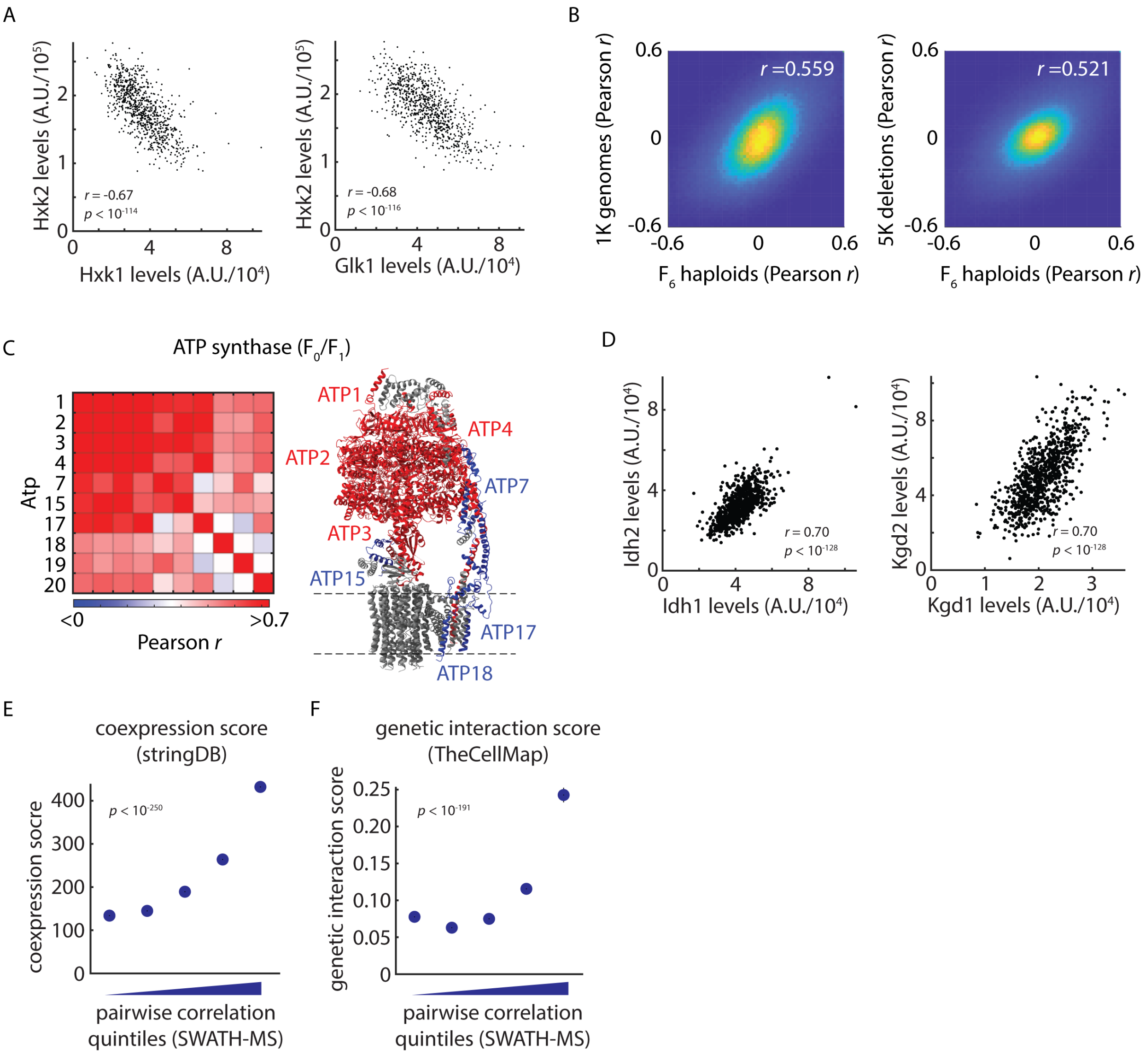


**Figure S5. To accompany Figure 5.** (A) Top: Hxk2 levels (ordinate) as a function of Hxk1 levels (abscissa) amongst $F_6$ progeny. Bottom: As above, but for Hxk2 and Glk1. (B) Left: Correlation between protein-protein correlations amongst 1,002 Genomes strains (ordinate) [32] and the same statistic amongst the $F_6$ progeny (abscissa). Shown is Pearson's *r*. Right: As on the left, for protein-protein correlations amongst precise deletions[30]. (C) Pairwise correlations of observed ATP synthase components (left) and cryo-EM structure of the yeast ATP synthase (6CP6)[86] with subunits highlighted as indicated (right). (D) Left: Idh2 levels (ordinate) as a function of Idh1 levels (abscissa) amongst $F_6$ progeny. Right: As left, but for Kgd2 and Kgd1. (E) Coexpression score from stringDB for ascending quintiles of protein pairs sorted by SWATH-MS abundance correlation. (F) As in (E) for the genetic interaction score from TheCellMap.

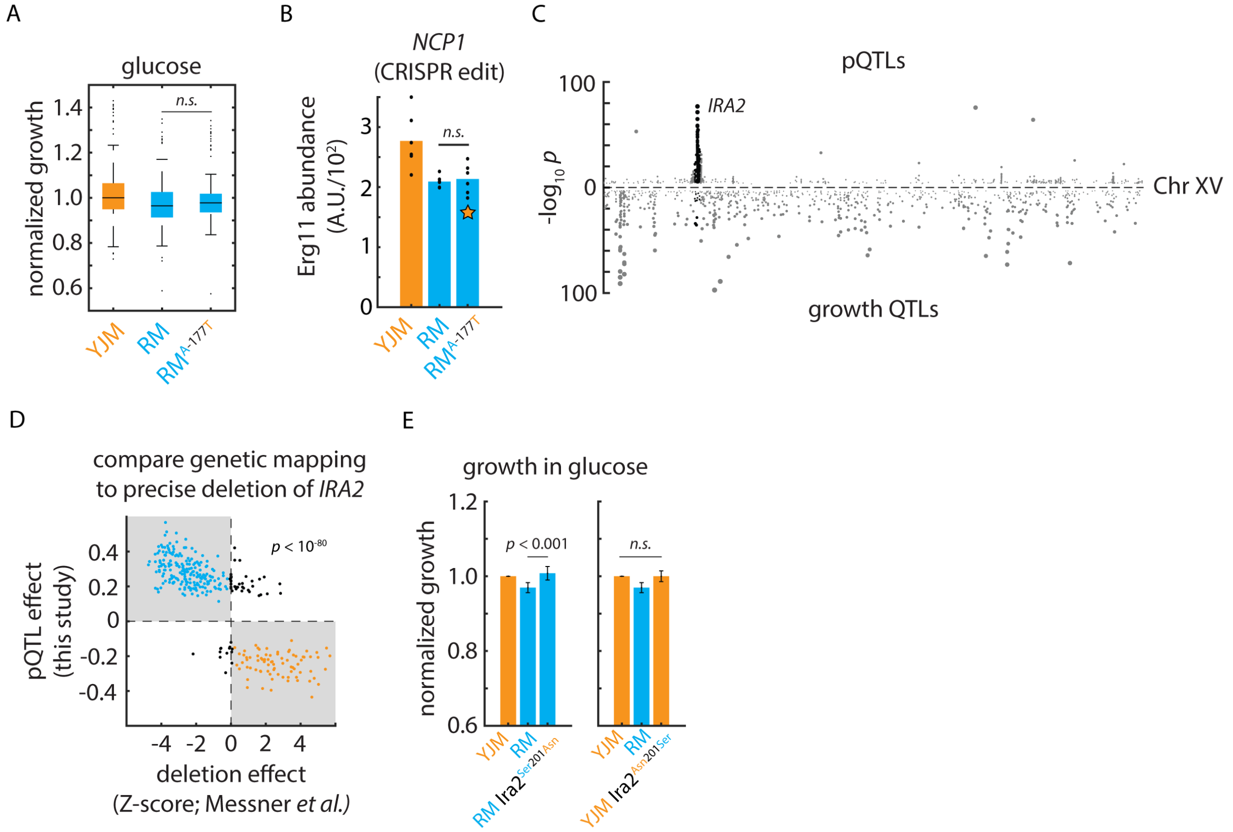


**Figure S6. To accompany Figure 6.** (A) Growth in glucose of clinical (YJM), vineyard (RM), and CRISPR-edited RM *NCP1*$^{A\text{-}177T}$ mutant strains in glucose. $n = 96$; $p$ value by Student's $t$ test. (B) CRISPR reconstruction and mass spectrometry to test the effect of the *NCP1*$^{\text{A-177T}}$ variant on Erg11 levels. $n = 6$; $p$ value by two-sided $t$ test. (C) Miami plot of pQTLs (top) and growth QTLs (bottom) on Chromosome XV. *IRA2* pQTLs and QTLs are highlighted in black. (D) Predicted *IRA2* pQTL effects from genetic mapping (this study; ordinate) as compared to measured effects of *IRA2* precise deletion (Z-scored by protein; [30]; abscissa) for all proteins predicted to be controlled by *IRA2*. $p$ value from $t$ statistic. Estimated abundances normalized to wild-type in each case. (E) Growth of clinical (YJM), vineyard (RM), and CRISPR-edited RM Ira2$^{\text{Ser201Asn}}$ mutant (left) and YJM Ira2$^{\text{Asn210Ser}}$ mutant (right) in glucose. $n = 96$; $p$ values by Student's $t$ test.

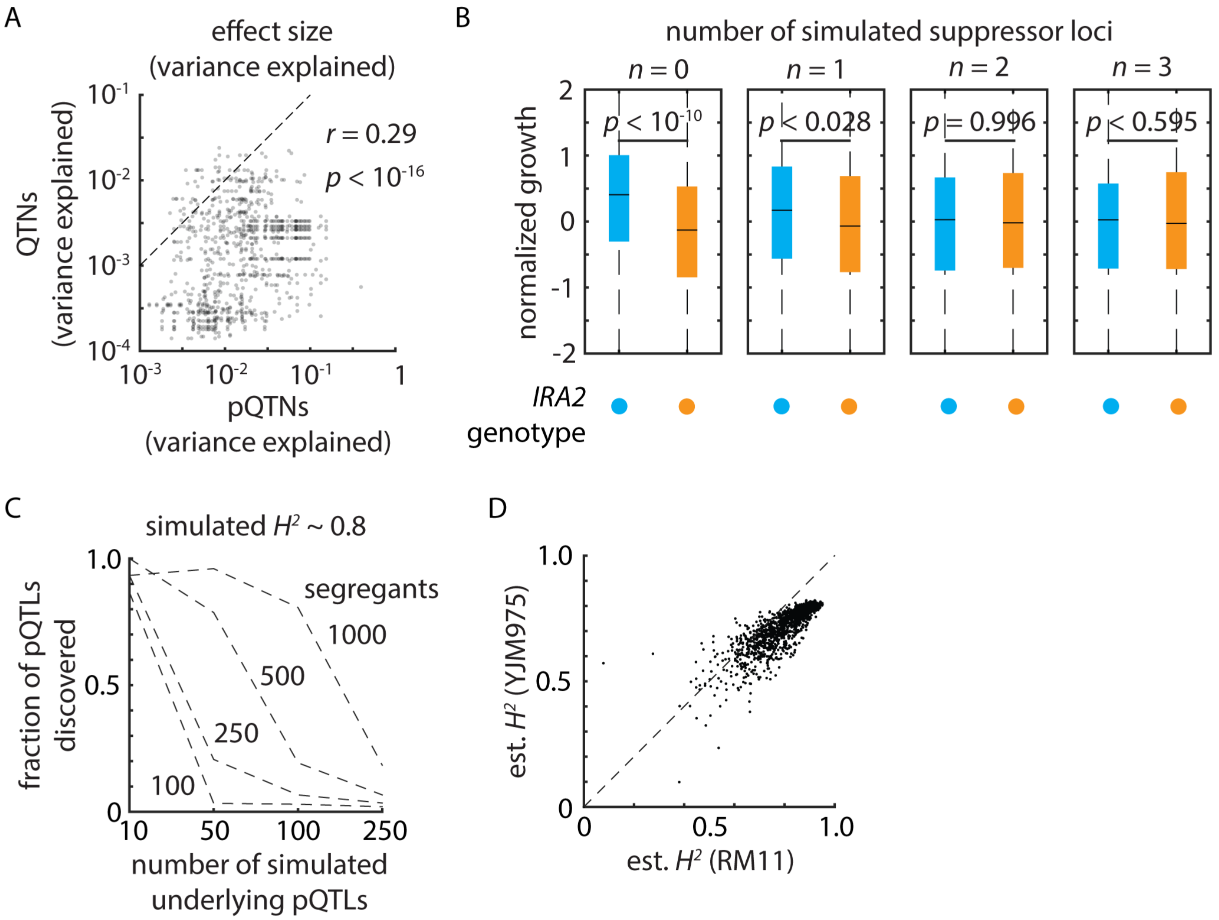


**Figure S7. To accompany Figure 7.**(A) Variance explained by phenotypic QTNs (ordinate) as a function of variance explained by pQTNs mapped to the same variant (abscissa). *r* by Pearson's correlation; *p* value from *t* statistic. (B) *In silico* simulations of the apparent effect of a linear QTL at *IRA2* with the number of segregating suppressing alleles indicated. *p* value by *t* test. (C) Fraction of simulated pQTLs discovered (ordinate) as a function of the number of true underlying simulated pQTLs (abscissa) for hypothetical pQTL mapping panels with the number of segregants indicated. N = 3 simulations for each parameter combination. (D) Estimated broad-sense heritability $H^2$ in YJM (ordinate) as a function of estimated $H^2$ in RM (abscissa) for all measured proteins.

| ***Upregulated in YJM975*** | | ***Upregulated in RM11*** | |
|---|---|---|---|
| **TF** | ***p* value** | **TF** | ***p* value** |
| Sfp1 | $< 10^{-22}$ | Sut1 | $< 10^{-11}$ |
| Stb3 | $< 10^{-16}$ | Msn2 | $< 10^{-11}$ |
| Abf1 | $< 10^{-10}$ | Msn4 | $< 10^{-11}$ |
| Sum1 | $< 10^{-10}$ | Hap3 | $< 10^{-9}$ |
| Gcn4 | $< 10^{-8}$ | Abf1 | $< 10^{-9}$ |
| Tod6 | $< 10^{-7}$ | Hap5 | $< 10^{-9}$ |
| Dot6 | $< 10^{-6}$ | Gis1 | $< 10^{-9}$ |
| Arg81 | $< 10^{-6}$ | | |
| Swi4 | $< 10^{-4}$ | | |

**Supplemental Table S6.** PSCAN transcription factor target enrichments and *p* values for proteins up- and down-regulated in the YJM975 and RM11 parents.
